# Effects of dopamine and serotonin on persistence for delayed rewards in humans

**DOI:** 10.64898/2026.08.11.744183

**Authors:** Karolina M. Lempert, Laura Zaneski, Arjun Ramakrishnan, Daniel H. Wolf, Joseph W. Kable

## Abstract

People often must decide how long to continue waiting for rewards that will arrive at an uncertain time in the future. We propose that these persistence decisions involve weighing the benefits of continued waiting against opportunity costs of waiting, a balance that may shift over time. This framework suggests that persistence decisions share neural mechanisms with foraging decisions, which require ongoing comparisons between a current resource and possible alternatives. Dopamine and serotonin have been proposed to play opposing roles in foraging, with dopamine promoting exploration and serotonin promoting exploitation. Here we investigated their roles in persistence. In a within-subjects, double-blind, placebo-controlled study in young adults (*n* = 42), we examined the effects of increasing dopamine with L-dopa and increasing serotonin with escitalopram. We predicted that L-dopa would decrease persistence and escitalopram would increase it. Participants also completed patch-foraging, time perception, risk tolerance, and temporal discounting tasks to explore potential mechanisms of drug effects on persistence. Escitalopram increased persistence, after adjusting for the effects of anxiety and condition order, such that participants waited longer for rewards after taking the serotonergic drug. L-dopa did not influence persistence. In exploratory analyses controlling for age, however, L-dopa reduced persistence and increased exploration in foraging.

## Introduction

People often face decisions about how long to wait for delayed rewards (i.e., *persistence decisions*). These decisions can concern people’s health (“how long should I stick with this diet if I haven’t seen results yet?”), finances (“how long should I hold onto this stock before selling?”), and even relationships (“how long should I continue to date this person if I haven’t yet felt a spark?”). Individual differences in how long people persist in waiting for delayed rewards are associated with real-world outcomes, with increased persistence associated positively with academic achievement and negatively with behavioral issues^1–3^. Research on these decisions also has considerable clinical relevance, given that many psychiatric disorders are marked by too much or too little persistence toward outcomes of value^4^. For example, individuals with obsessive-compulsive disorder are unable to stop performing elaborate rituals that they believe will make them feel better despite the opportunities they lose by doing so (over-persistence). On the other hand, individuals with substance use disorder have difficulty persisting toward the sobriety they desire (under-persistence). A better understanding of the cognitive and neural mechanisms of persistence decisions will potentially open up new avenues for treatment. In the current study, we tested the novel hypothesis that dopamine and serotonin have opponent roles in persistence for delayed rewards, by having healthy young adults take single doses of levodopa (L-dopa; a dopamine precursor that increases dopamine levels) and escitalopram (a selective serotonin reuptake inhibitor, or SSRI, that increases serotonin levels) prior to performing a well-validated persistence task.

In persistence decisions, people frequently quit waiting for delayed rewards even after an initial decision to wait. Such “changes of mind” have often been interpreted as irrational^5,6^: if it was not worth waiting the full time until the reward arrived, then one should not have chosen to wait initially. However, there is often uncertainty about when delayed rewards will arrive. Given this temporal uncertainty, quitting can be the rational choice if a person’s beliefs about how much longer they have to wait change over time^7^. That is, the choice whether to continue waiting or to quit may be the result of *dynamic reassessments* during the waiting period. For instance, you might start a diet because you expect to see results in a week; if you do not, then you might quit the diet because you *now* believe that it will take much longer to work, and so it is not worth forgoing delicious foods any longer. Under this dynamic reassessment hypothesis, how long people wait for delayed rewards reflects their (1) expectations about the possible times at which delayed rewards will arrive, and (2) perceptions of the *opportunity cost* of persisting (i.e., rewards they miss out on by waiting). This conceptualization of persistence differs from theories that presuppose that people are motivated to wait for the delayed reward throughout the delay period, but their persistence is curtailed due to a failure of behavioral inhibition or willpower^8,9^.

When conceptualized as dynamic, persistence structurally resembles foraging, which has been studied extensively in humans^10–13^ and non-human animals^14–17^. In foraging, animals dynamically decide whether to continue exploiting a current resource (e.g., a tree or bush) or to give up and explore other, potentially richer, options. By borrowing insights from foraging theory, one can model, during persistence, the value of waiting relative to quitting at different points in time, given beliefs about reward timing^18,19^. When foraging, according to the Marginal Value Theorem^20,21^, an animal should give up on the current resource when the average reward rate in the environment – across all alternative resource options – exceeds the reward rate of the current resource. Analogously, when waiting for delayed rewards, one should choose to quit waiting for a reward when the opportunity cost of waiting exceeds the expected value of the current reward, given beliefs about its timing.

This account of persistence decisions is supported by previous research that used a persistence task similar to the one we used here. The goal of the task is to collect as many small (10¢) rewards as possible in a given amount of time. In each trial, participants wait through a variable delay for the small reward. At any time in the delay period, participants can choose to quit waiting and move on to a different trial. The delay length is unknown to participants, and it can be drawn from different distributions that dictate different optimal waiting times based on the foraging-inspired model. In these studies, participants are able to learn reward timing statistics from experience, and to adjust how long they wait accordingly, waiting less when waiting is costly and waiting longer when it pays off^18,22,23^. They successfully trade off the benefits of waiting against the opportunities lost by waiting. Thus, in both foraging and persistence, decision makers weight the reward expected from the current resource against their opportunity costs, or what they would expect from exploring other options.

Two neurotransmitters – dopamine and serotonin – are proposed to play important and opposing roles in foraging decisions^24–28^, with dopamine promoting exploration and serotonin promoting exploitation. Here we probed the role of these neurotransmitters in persistence decisions, by having participants perform a persistence task three times: once on placebo, once on escitalopram, and once on L-dopa. Although the mechanisms of action of escitalopram and L-dopa differ – escitalopram is a selective serotonin reuptake inhibitor, whereas L-dopa is a precursor to dopamine – they increase tonic levels of serotonin and dopamine, respectively.

Important theoretical work proposes that tonic dopamine levels in the brain signal the average reward rate in an environment^29^. This average reward rate, across all resource alternatives, represents the opportunity cost of exploiting a single current resource. This theory predicts that an increase in tonic dopamine would lead to more exploratory foraging choices (i.e., leaving a resource sooner), by increasing perceived opportunity cost. Pharmacological manipulations of dopamine in humans lead to changes in decision making in foraging-like tasks, but these effects vary. In one study^27^, individuals with Parkinson’s disease made more exploratory foraging choices when they were on L-dopa compared to when they were off medication, an effect that was not explained by motor perseveration, and that did not differ as a function of the average reward rate of the environment. However, other studies have suggested that dopamine, rather than signaling average reward rate directly, alters the extent to which average reward rate modulates decision making. For example, healthy young adults given L-dopa made faster, more vigorous responses as the average reward rate of an environment increased^28^. On the other hand, in older adults, the D2 receptor agonist cabergoline *reduced* the effect of environment reward rate on decisions, such that participants were more likely to explore even when environment reward rates were low^30^. In line with this, a D2 receptor antagonist, amisulpride, has been shown to increase the effect of environment reward rate on foraging decisions^31^. Despite these mixed findings, given the parallels between foraging and persistence, and the theory that tonic dopamine signals average reward rate, we expected that an increase in tonic dopamine would lead to less willingness to wait through a delay, since the opportunity cost of waiting (i.e., the average reward rate of the environment) would be perceived as higher. Thus, we predicted that L-dopa would lead participants to quit waiting for delayed rewards sooner (Hypothesis 1). Alternatively, L-dopa might lead to no change in persistence (H1_0_). See Table 1 for the full list of hypotheses.

**Table 1.** Design Table.

| Hypotheses | Outcome Measures | Sampling Plan (N, power analyses) | Analysis Plan | Potential outcomes & interpretations | Result & interpretation |
| --- | --- | --- | --- | --- | --- |
| *H1<sub>1</sub>*: Persistence will be reduced on L-dopa compared to placebo. | Area under the curve (AUC) in persistence task | Planned $n = 42$ participants, based on medium effect size Cohen's $d_z = 0.425$, power = 94% | Mixed-effects linear regression on subset of the data, examining effects of medication condition (Placebo vs. L-dopa) on AUC with potential covariates (see Supplementary Table 1), with random intercepts and slopes on medication condition by subject. | *Outcome 1.* Significantly ($p < 0.025$) lower AUC in L-dopa condition compared to placebo: Evidence for H1.<br>*Outcome 2.* No significant difference between conditions ($p > 0.025$): Lack of evidence for H1.<br>*Outcome 3.* Significantly ($p < 0.025$) higher AUC in L-dopa condition compared to placebo: Lack of evidence for H1. | *Outcome 2.* No significant difference between conditions ($\beta = -0.89$; $p = 0.841$). Lack of evidence for H1. |
| *H1<sub>0</sub>*: Persistence in L-dopa and placebo conditions will be statistically equivalent. | Area under the curve (AUC) in persistence task | Planned $n = 42$ participants, based on medium effect size Cohen's $d_z = 0.425$, power = 94% | Two one-sided $t$-test (TOST) procedure, assuming smallest effect size of interest Cohen's $d_z = 0.14$. | *Outcome 1.* Both one-sided $t$-tests (TOST upper and TOST lower) are significant ($p < 0.05$): Evidence for statistical equivalence (H1<sub>0</sub>).<br>*Outcome 2.* Only one $t$-test (TOST upper or TOST lower) is significant ($p < 0.05$): Lack of evidence for H1<sub>0</sub>.<br>*Outcome 3.* Both one-sided $t$-tests are not significant ($p > 0.05$): Lack of evidence for H1<sub>0</sub>. | *Outcome 2.* TOST lower test was significant ($t_{41} = 2.06$; $p = 0.023$), but TOST upper test was not ($t_{41} = 0.25$; $p = 0.598$). Lack of evidence for H1<sub>0</sub>. |
| *H2<sub>1</sub>*: Persistence will be increased on escitalopram compared to placebo. | Area under the curve (AUC) in persistence task | Planned $n = 42$ participants, based on medium effect size Cohen's $d_z = 0.425$, power = 94% | Mixed-effects linear regression on subset of the data, examining effects of medication condition (Placebo vs. Escitalopram) on AUC with potential covariates (see Supplementary Table 1), with random intercepts and slopes on medication condition by subject | *Outcome 1.* Significantly ($p < 0.025$) higher AUC in escitalopram condition compared to placebo: Evidence for H2.<br>*Outcome 2.* No significant difference between conditions ($p > 0.025$): Lack of evidence for H2.<br>*Outcome 3.* Significantly ($p < 0.025$) lower AUC in escitalopram condition compared to placebo: Lack of evidence for H2. | *Outcome 1.* Significantly higher AUC in escitalopram condition compared to placebo ($\beta = 3.92$; $p = 0.012$). Evidence for H2. |
| *H2<sub>0</sub>*: Persistence in escitalopram and placebo conditions will be statistically equivalent. | Area under the curve (AUC) in persistence task | Planned $n = 42$ participants, based on medium effect size Cohen's $d_z = 0.425$, power = 94% | Two one-sided $t$-test (TOST) procedure, assuming smallest effect size of interest Cohen's $d_z = 0.14$. | *Outcome 1.* Both one-sided $t$-tests (TOST upper and TOST lower) are significant ($p < 0.05$): Evidence for statistical equivalence (H2<sub>0</sub>).<br>*Outcome 2.* Only one $t$-test (TOST upper or TOST lower) is significant ($p < 0.05$): Lack of evidence for H2<sub>0</sub>.<br>*Outcome 3.* Both one-sided $t$-tests are not significant ($p > 0.05$): Lack of evidence for H2<sub>0</sub>. | Test is not applicable.<br>H2 is supported; H2<sub>0</sub> is rejected. |

Serotonin is proposed to play an opposing role to dopamine in foraging. Specifically, rather than signaling the average reward rate, serotonin may signal the average punishment rate of the environment^32,33^. Assuming a strict opponent relationship between rewards and punishments (which may not necessarily be the case; for discussion, see^34^), then in the absence of aversive feedback, the average punishment rate is equivalent to the inverse of the average reward rate. Therefore, an increase in tonic serotonin should lead an animal to *exploit* the current option because the cost of exploring is high. Consistent with this idea, serotonin plays a role in inhibiting exploration and promoting exploitation in non-human animals. The dorsal raphe nucleus, a key source of serotonin in the brain, has been shown to respond to the average reward rate of the environment during decision making in macaque monkeys^35,36^. Acute administration of citalopram, an SSRI, increased the amount of time macaque monkeys waited to act during a persistence task^36^. In mice, serotonin neurons fire more during waiting for rewards^25^. Inhibiting these neurons promotes exploration^37^, and optogenetic activation of these neurons promotes waiting^26,38–40^. Serotonin does not inhibit actions indiscriminately, however; in one study, optogenetic activation of serotonin neurons increased the exploitation of a current resource, even when actions had to be taken to exploit it^24^. This research in non-human animals supports the prediction that an increase in tonic serotonin should promote persistence for delayed rewards. However, just like with dopamine, some research suggests that serotonin does not always increase patience or exploitation, but that it modulates the influence of background reward rate on choice. Indeed, disrupting the dorsal raphe nucleus diminishes an animal’s sensitivity to the reward environment^36,41^. Notably, however, dorsal raphe nucleus activity specifically encoded transitions from rich to poor environments, in which an animal switches from a vigorous, exploratory mode to a slower, more exploitative mode^41^. Here we increased postsynaptic serotonin levels acutely with the SSRI escitalopram^42^. In our persistence task, we expected that escitalopram would lead participants to wait longer for delayed rewards (Hypothesis 2). Alternatively, escitalopram might lead to no change in persistence (H2_0_). In fact, one study that manipulated the average reward rate during a discrimination task in humans found that an SSRI had no effect on motor vigor as the average reward rate decreased^28^. This null result contradicts a central prediction of the serotonin/dopamine opponency theory^33,34^. One possible explanation for this result is that serotonin might only elicit exploitation and behavioral inhibition in contexts that involve actual punishment (e.g., monetary loss, shock)^43–45^. Our investigation helps address this important question.

In the persistence task that we used here, reward times were drawn from a quasi-exponential distribution, such that participants would earn approximately the same amount of money regardless of how long they chose to wait (Supplementary Fig. 1). Thus, we did not externally manipulate opportunity cost by varying the reward timing distributions that participants encountered. Rather, we predicted that differences between medication conditions would stem from *endogenous* changes in perceptions of opportunity cost, or of the value of the awaited reward. We made this choice because previous research has shown that people learn on the basis of feedback how to shift their wait times in response to changing opportunity costs in this task,^46^ and we did not want differences in behavior between medication conditions to be potentially driven by differences in a participant’s ability to learn the statistics of a timing environment. For example, if the optimal waiting policy was to wait only a short time before quitting, then shorter wait times might reflect enhanced learning of reward timing statistics rather than increased exploration. Thus, in the current task, we can only test the hypotheses that dopamine and serotonin signal average reward rate directly; we cannot examine whether the medications influence how people modulate their behavior in response to external manipulations of opportunity cost.

If we were to find support for one or both of our hypotheses, we wanted to establish the mechanism by which these medications influence persistence. To this end, we also measured the effects of L-dopa and escitalopram on behavior in several other tasks. One possibility is that dopamine and serotonin may affect persistence by altering perceptions of opportunity cost. To test this mechanism, we included a standard patch-foraging task^13,16,17^, in which subjects collect rewards from patches and must decide when to move on from each patch as it gets depleted. In the L-dopa condition, if increasing dopamine transmission increases perceived opportunity cost, subjects should leave patches earlier. In contrast, if increasing serotonin transmission decreases perceived opportunity cost, escitalopram should lead participants to stay longer in patches. This pattern of results, if found, would support an opportunity cost mechanism for the persistence task results, and we would do a formal mediation analysis to confirm this mechanism. It is also possible that the results of the foraging and persistence tasks might diverge. Indeed, many theoretical accounts of persistence under temporal uncertainty (for discussion, see^7^) do not draw an analogy between willingness-to-wait and foraging. The current study sheds light on the extent to which these two processes engage similar neural mechanisms. An added benefit of including a foraging task is the opportunity to replicate previous pharmacological studies on foraging^27,30^ in healthy young adults.

We also planned to test other possible mechanisms by which dopamine and serotonin might influence persistence: specifically, through their effects on temporal discounting, time perception, and/or risk tolerance. Temporal discounting refers to the tendency to prefer rewards sooner rather than later, and is typically measured by having participants make decisions between smaller, sooner monetary rewards and larger rewards that are delayed by some number of days (e.g., “$10 now vs. $20 in 7 days”). Higher temporal discounting rates correspond to more impatience. We view temporal discounting as distinct from persistence for delayed rewards under uncertainty. In a temporal discounting task, choosing delayed rewards (1) does not entail temporal uncertainty, since the delay is clearly indicated, and (2) does not give rise to opportunity costs, since subjects can do other activities during the days-long waiting periods. Of course, it is possible that a change in temporal discounting between conditions might drive changes in persistence between conditions (e.g., if both tasks capture an aversion to waiting for rewards in general). Therefore, we measured temporal discounting in each medication condition. Temporal discounting has been linked with both dopamine and serotonin transmission, although these findings are mixed. Regarding serotonin, one study found that serotonin depletion in humans increased temporal discounting^47^, while others found null effects^48,49^. Regarding dopamine, one study found that L-dopa increased temporal discounting^50^, others found that it decreased temporal discounting (in healthy participants^51^ and in individuals with Parkinson’s disease^52^), and yet others found no effect^53^. Studies that administered D2-receptor antagonists – which may in some doses actually increase presynaptic dopamine transmission^54^ – found that they decreased temporal discounting^55–57^ or had no effect^50^. These heterogeneous findings, combined with the lack of an association between dopamine receptor availability and temporal discounting in healthy adults^58^, suggest that perhaps dopamine does not affect temporal discounting directly, but rather may only influence these decisions through other means (e.g., via effects on general reward processing^59^). If we were to find changes in temporal discounting between medication conditions, however, we planned to test whether medication effects on persistence are mediated through temporal discounting.

We also examined effects of L-dopa and escitalopram on time perception and risk tolerance. Accurately judging the lengths of time intervals can influence persistence^60–62^, and can be affected by dopamine transmission^63^. Participants therefore completed a time production task^64^ in each medication condition. L-dopa and/or escitalopram may also influence risk tolerance^65,66^, which might then affect persistence. Risk tolerance was assessed with a standard risky choice task (e.g., “$20 for sure vs. $50 with 50% chance”) at each test visit. If we were to find any significant effects of medication on time perception or risk tolerance, we planned to conduct mediation analyses to examine whether our persistence results were driven by time perception or risk tolerance.

In summary, here in a within-subjects, double-blind, placebo-controlled study, we tested the hypotheses that an increase in tonic dopamine levels due to L-dopa would decrease persistence for delayed rewards (Hyp. 1), and that an increase in serotonin levels due to escitalopram would increase persistence for delayed rewards (Hyp. 2). Contingent on finding support for these hypotheses, we also planned to conduct exploratory analyses of secondary tasks, in order to constrain our interpretation of the results. If these medications affect persistence by altering perceived opportunity cost, then we would expect to observe decreased patch staying times on L-dopa and increased patch staying times on escitalopram in a foraging task. We did not expect that our results would be driven by changes in time perception, temporal discounting, or risk tolerance, but we included those measures in order to test those alternative mechanisms.

## Results

### Participants

Forty-two participants (mean age = 26.8; SD = 7.40; range: 18-44; 25 F, 17 M; 15 Asian, 10 White, 5 Black, 5 Hispanic or Latino, 1 American Indian, 1 mixed race, 5 other race or not reported) passed a health screening and completed three test visits. At each test visit, they received either a placebo, 20 mg of escitalopram (“SSRI”), or 200 mg of levodopa + 50 mg of carbidopa (“L-dopa”). The condition order was counterbalanced across participants. Once the medications were close to their peak concentrations (3 hours after SSRI, and 90 minutes after L-dopa), participants then performed a series of decision-making tasks, including a persistence task (see *Methods*; Figs. 1-2). There were no serious adverse events, so all participants who completed the study were included in the final analysis. However, four participants experienced nausea, two of whom also vomited. Three of those participants had taken the SSRI, and one had taken L-dopa. One additional participant experienced dizziness after taking the SSRI.

**Fig. 1.**
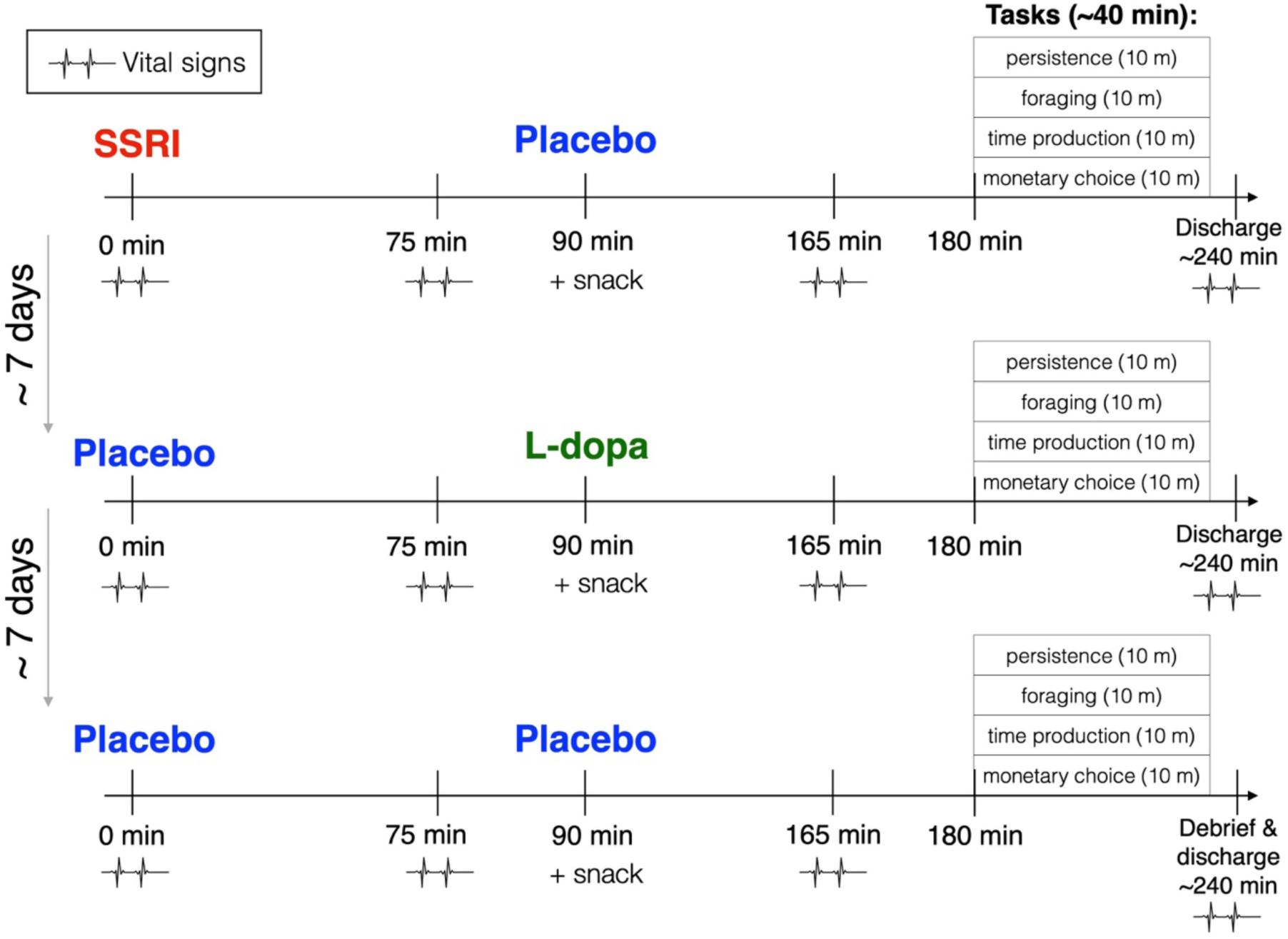
Diagram of procedure for test visits. Example procedure for a participant, for the example order of SSRI-L-dopa-Placebo. Each row corresponds to a visit. The visits differed only with respect to which medications were given. The test visit began with medication (or placebo) administration, followed by a 1.5-hour waiting period until a snack and a second round of medication (or placebo) administration. During the waiting period on the first day, subjects completed baseline questionnaire and task measures. After 3 hours, the participant completed the four experimental tasks. While the tasks were done in the same order at each visit, and the persistence task was presented first, the order of the remaining three tasks (foraging, time production, and monetary choice) was counterbalanced across subjects. Vital signs were assessed approximately each hour for safety.

**Fig. 2.**
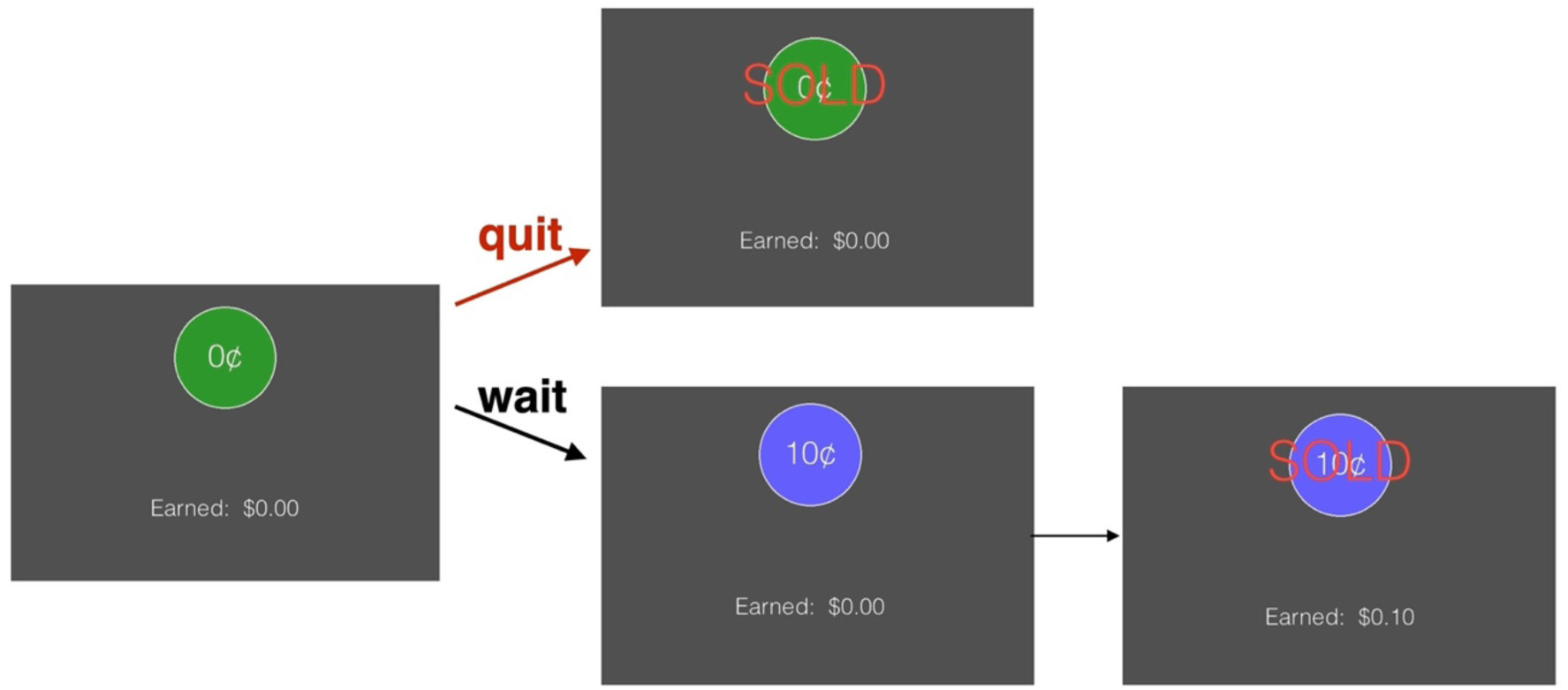
One trial of the persistence task. Participants can either wait for the token to mature to sell it, or sell the token before it matures to advance to the next trial. Once they press the spacebar to sell the token, the word “SOLD” appears over the token for 0.25 s, and then a new trial appears on the screen. Accumulated earnings are shown on the bottom of the screen.

### Pre-registered Analyses

#### Blinding was effective

To test whether participants were aware of which medication they were given, we asked them to guess the order in which they received the three treatments (Placebo, L-dopa, and SSRI) at the end of the study. We performed a one-sample *t*-test to compare the proportion of correct guesses to chance level performance, which is 1/6 (16.67%). Four participants were excluded from this analysis for guessing “placebo” twice. Out of the remaining 38 participants, only six guessed the order correctly, suggesting that blinding was successful (*t*_37_ = -0.15; *p* = 0.884). Next, we tested if participants were above chance in guessing whether or not they had taken a placebo versus an active medication. Only 26.3% of participants guessed the placebo day correctly, a proportion which was no different from chance (33.33%), again suggesting that blinding was successful (*t*_37_ = -0.97; *p* = 0.339).

#### No effect of L-dopa on persistence

We predicted that L-dopa would reduce persistence, measured by AUC (see *Methods*). Prior to testing this hypothesis (Hyp. 1), we first examined if there were any covariates that we needed to adjust for in our main model. We started by testing if body mass index (BMI), age, sex (Female / Male), condition order (Placebo first / L-dopa first), working memory capacity (OSPAN partial score), or Barratt Impulsiveness Scale (BIS) score was significantly associated with AUC, or if any of these covariates interacted with L-dopa to influence AUC. To this end, we constructed a series of mixed-effects linear regression models. None of the covariates yielded significant effects (all *p*s > 0.05), with the exception of the BIS score, which significantly predicted AUC (β = -0.23; *p* = 0.049). We also tested to see if there was a main effect of L-dopa on positive affect (from the Positive and Negative Affect Schedule, PANAS-P), negative affect (PANAS-N) or state anxiety (as assessed with the Spielberger Trait Anxiety Inventory – State version; STAI-S), or if L-dopa interacted with timepoint (before vs. after medication) to predict any of these variables. There were no effects of L-dopa on any of these affective variables (all *p*s > 0.05).

Therefore, the final model testing for an effect of L-dopa on AUC included the following predictors: L-dopa, BIS score, and L-dopa x BIS score. Random effects were included for both L-dopa and BIS score, but not their interaction term (due to nonconvergence of the more complex model). In this model, there was no main effect of L-dopa on persistence (β = -0.89; 95% CI: [-9.58, 7.80]; *p* = 0.841; Table 2; Fig. 3).

**Fig. 3.**
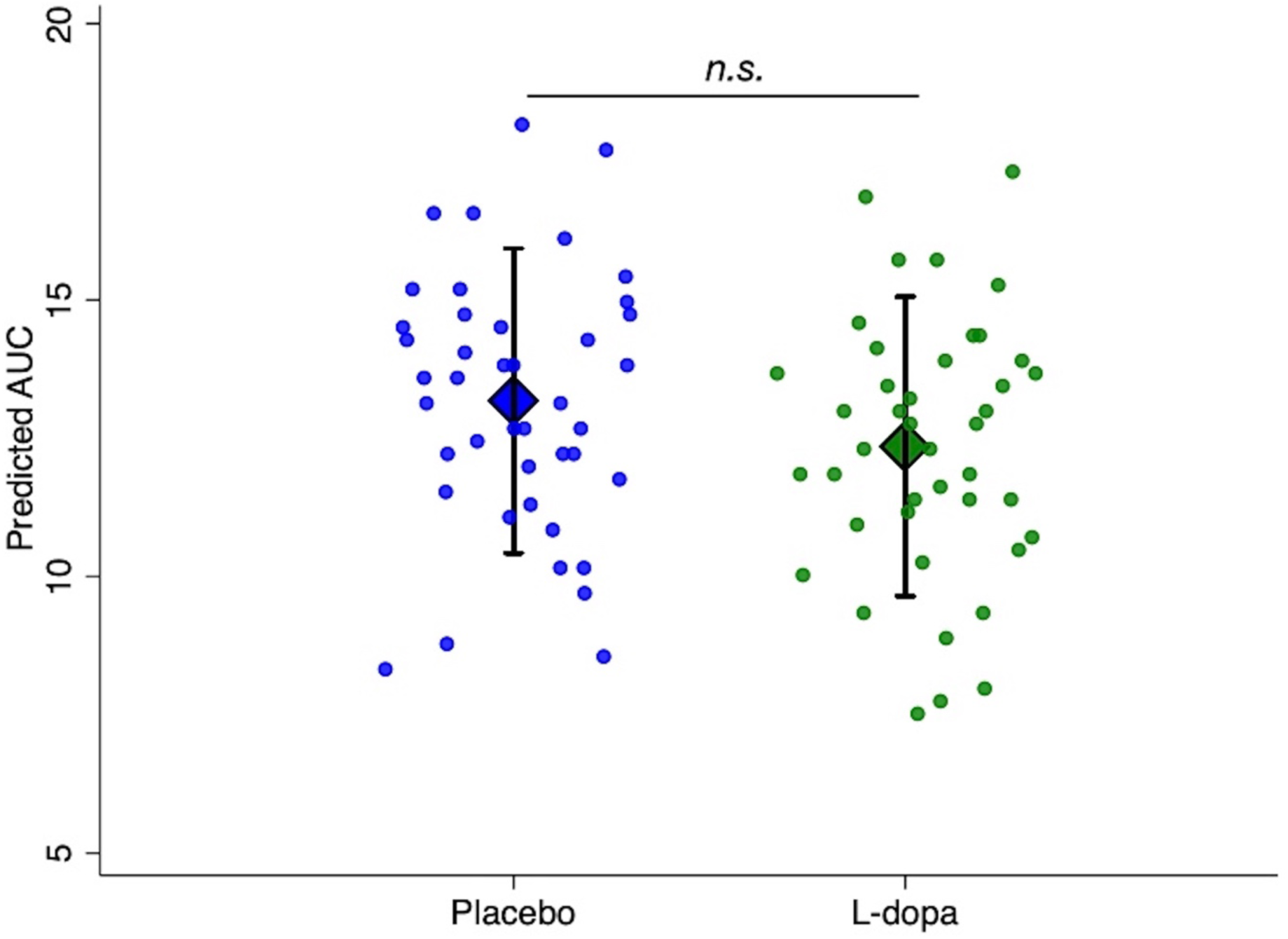
Estimated marginal means for persistence for placebo and L-dopa conditions, with individual adjusted data points. Diamond markers show model-estimated marginal means of area under the curve (AUC) from a mixed-effects regression including medication and Barratt Impulsiveness Scale (BIS) scores, with error bars indicating 95% confidence intervals. Overlaid points represent individual participants’ AUC residualized with respect to BIS scores, plotted with horizontal jitter for visibility.

**Table 2.** Fixed effects of L-dopa on persistence (AUC)

| Predictor | $\beta$ | 95% CI |
| --- | --- | --- |
| L-dopa<br>(1 = L-dopa; 0 = Placebo) | -0.89 | -9.58, 7.80 |
| BIS score | -0.23* | -0.46, -0.001 |
| L-Dopa x BIS score | 0.001 | -0.14, 0.14 |

Note: Dependent variable is area under the curve (AUC) from persistence task, with higher values indicating longer wait times. Random effects were included for the intercept and L-dopa terms. CI = confidence interval. BIS = Barratt Impulsiveness Scale. \* $p < 0.05$ .

Since there was no significant effect of L-dopa on persistence, we next conducted a TOST test for statistical equivalence, assuming a smallest effect size of interest of Cohen’s *d_z_* = 0.14 (for this comparison, Cohen’s d_z_ = -0.18; 95% CI: [-0.48, 0.13]). Whereas one of the one-sided *t*-tests was significant (TOST lower, *t*_41_ = 2.06; *p* = 0.023), the other was not (TOST upper, *t*41 = 0.25; *p* = 0.598). Therefore, although persistence in the L-dopa condition was not statistically different from the placebo condition, we do not have sufficient evidence to conclude that behavior in the two conditions was statistically equivalent at our chosen threshold. Therefore, we cannot reject the null hypothesis Hyp. 1_0_.

#### SSRI increases persistence

Next, we tested whether the SSRI escitalopram would increase persistence (Hyp. 2). Once again, we first performed analyses to determine if BMI, age, sex, or condition order influenced AUC, or if any of these variables interacted with SSRI to predict AUC. We found that there was a significant interaction between SSRI and condition order (Placebo first / SSRI first) on AUC (β = −4.31; *p* = 0.032). When examining medication effects on positive affect, negative affect, and state anxiety, we found a significant interaction between SSRI and timepoint (before vs. after medication) on state anxiety (β = 3.55; *p* = 0.042). Follow-up paired-samples *t*-tests revealed that this interaction was driven by an increase in anxiety in the SSRI condition (*t*_41_ = −2.80; *p* = 0.008) and no change in anxiety in the placebo condition (*t*_41_ = 0.12; *p* = 0.909).

Therefore, the final model testing for an effect of SSRI on AUC contained the SSRI predictor, a condition order covariate (Placebo first / SSRI first), a change in state anxiety covariate (STAI-S after – STAI-S before = ΔSTAI), as well as all two-way and three-way interaction terms among predictors. In this model, there was a significant positive effect of SSRI on persistence (β = 3.92; *p* = 0.012; Table 3). In addition, there was a significant interaction between SSRI and condition order on persistence (β = -5.53; *p* = 0.011), as well as a significant interaction between SSRI and ΔSTAI on persistence (β = -0.47; *p* = 0.016). All other effects in the model were not significant (all *p*s > 0.05). Therefore, taking the SSRI increased the length of time that people were willing to wait for delayed rewards. Hyp. 2 was supported and the null hypothesis Hyp. 2_0_ was rejected. However, this effect was qualified by two interactions.

**Table 3.** Fixed effects of SSRI on persistence (AUC).

| Predictor | $\beta$ | 95% CI |
| --- | --- | --- |
| SSRI<br>(1 = SSRI; 0 = Placebo) | 3.93* | 0.87, 6.98 |
| Condition order<br>(1 = Placebo first; 0 = SSRI first) | -0.86 | -6.53, 4.81 |
| SSRI x Condition order | -5.53* | -9.78, -1.27 |
| ΔSTAI-S | 0.18 | -0.15, 0.51 |
| SSRI x ΔSTAI-S | -0.47* | -0.86, -0.09 |
| Condition order x ΔSTAI-S | -0.37 | -0.86, 0.12 |
| SSRI x ΔSTAI-S x Condition order | 0.59 | -0.10, 1.29 |

Note: Dependent variable is area under the curve (AUC) from persistence task, with higher values indicating longer wait times. Random effects were included for the intercept and SSRI terms. CI = confidence interval. SSRI = selective serotonin reuptake inhibitor (20 mg escitalopram). ΔSTAI-S = change in Spielberger State Anxiety Inventory score (after medication – before medication). \* $p < 0.05$ .

First, the effect of the SSRI on persistence was stronger when the SSRI test visit preceded the placebo test visit (Fig. 4). Estimated marginal means were computed for each condition order and medication condition combination, followed by pairwise comparisons. Pairwise comparisons between the SSRI and placebo conditions showed a significant difference in AUC when the SSRI was given first (mean difference between SSRI+SSRI first and Placebo+SSRI first = 3.15; 95% CI [0.29-6.01], *p* = 0.031), but not when the SSRI was given after placebo (mean difference between SSRI+Placebo first and Placebo+Placebo first = -1.41; 95% CI [-4.49-1.67], *p* = 0.370). Moreover, when participants had taken the SSRI, they were more willing to wait if the SSRI visit preceded the Placebo visit (mean difference between SSRI+SSRI first and SSRI+Placebo first = 6.02; 95% CI [0.65-11.39], *p* = 0.028). On the other hand, when participants had taken the placebo, their persistence did not depend on condition order (mean difference between Placebo+SSRI first and Placebo+Placebo first = 1.46; 95% CI [-4.29-7.21], *p* = 0.618). Therefore, the effects of the serotonergic medication on persistence were weaker when participants had already been exposed to the task. Indeed, regardless of the order in which they were given the medications, participants’ AUCs tended to decrease from the first test visit to the second test visit (mean AUC at Test Visit 1 = 14.4; mean AUC at Test Visit 2 = 12.4; *t*_41_ = 2.23; *p* = 0.031), before stabilizing (mean AUC at Test Visit 3 = 12.2; comparison to Test Visit 2: *t*_41_ = 0.21; *p* = 0.833).

**Fig. 4.**
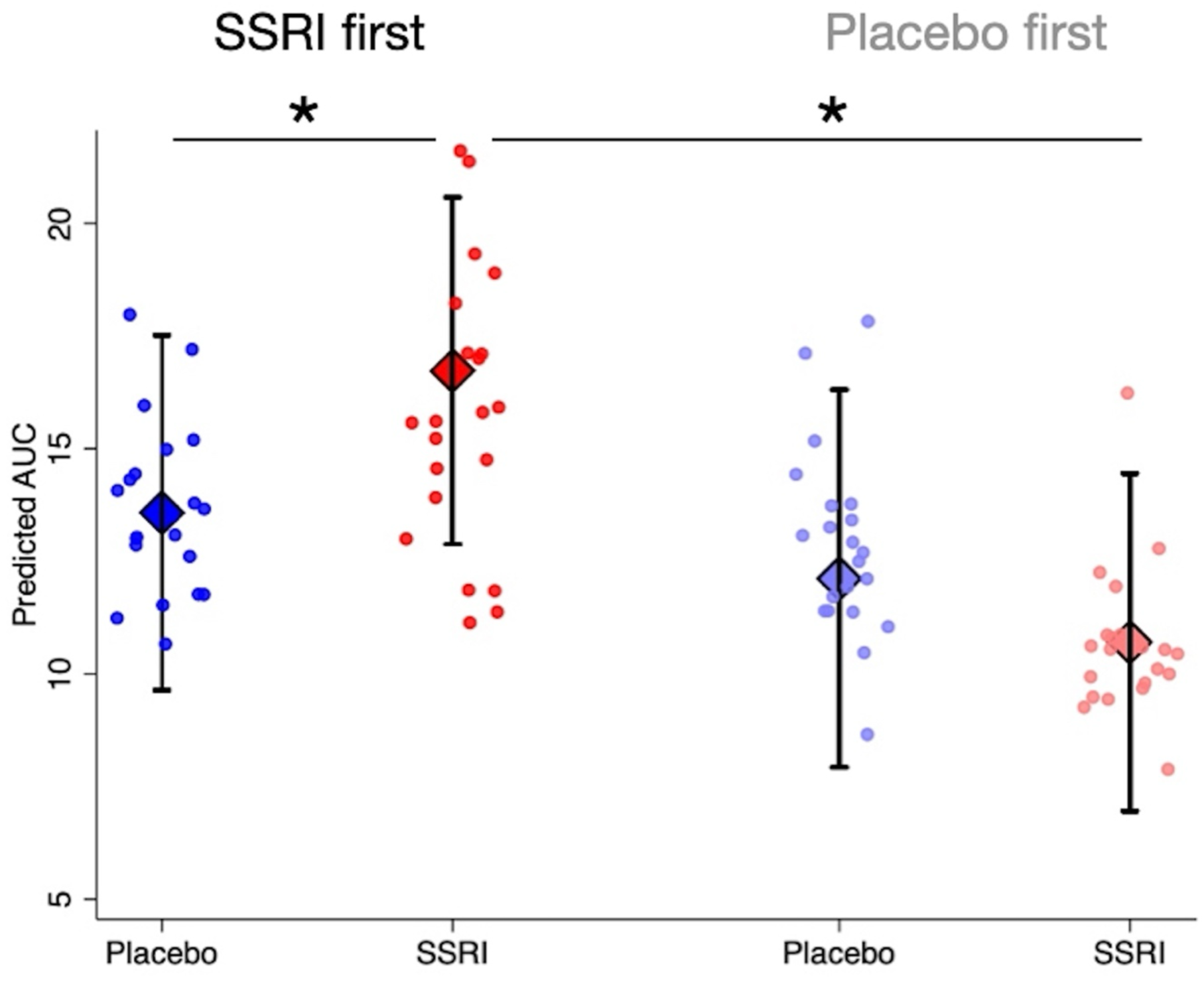
Estimated marginal means for persistence, by medication condition and condition order, with individual adjusted data points. Diamond markers show model-estimated marginal means of area under the curve (AUC) from a mixed-effects regression including medication, condition order, and change in state anxiety, with error bars indicating 95% confidence intervals. Darker diamonds and points correspond to when the selective serotonin reuptake inhibitor (SSRI) was given before placebo, and lighter diamonds and points correspond to when placebo was given before the SSRI. Overlaid points represent individual participants’ AUC residualized with respect to change in state anxiety, plotted with horizontal jitter for visibility. \**p*<0.05.

Second, in the SSRI condition, an increase in anxiety was associated with a reduction in persistence (Fig. 5). To understand what was driving the interaction between anxiety change (ΔSTAI) and medication, we calculated the simple slopes between ΔSTAI and AUC, for each combination of medication condition and condition order (note: although the three-way interaction among ΔSTAI, condition order and medication condition did not reach significance, the two-way interaction between ΔSTAI and medication condition was significant only when condition order was in the model). We found that there was a significant relationship between anxiety change and persistence when participants were in the SSRI condition, and they took the SSRI prior to taking placebo (SSRI+SSRI first: β = -0.29; 95% CI [-0.57, -0.02], *p* = 0.037). All other slopes were not significantly different from zero (all *p*s > 0.05). Therefore, while taking the SSRI led to an increase in persistence (particularly if the SSRI session was prior to the placebo session), participants who were the most anxious after taking the SSRI were the least likely to show the effect.

**Fig. 5.**
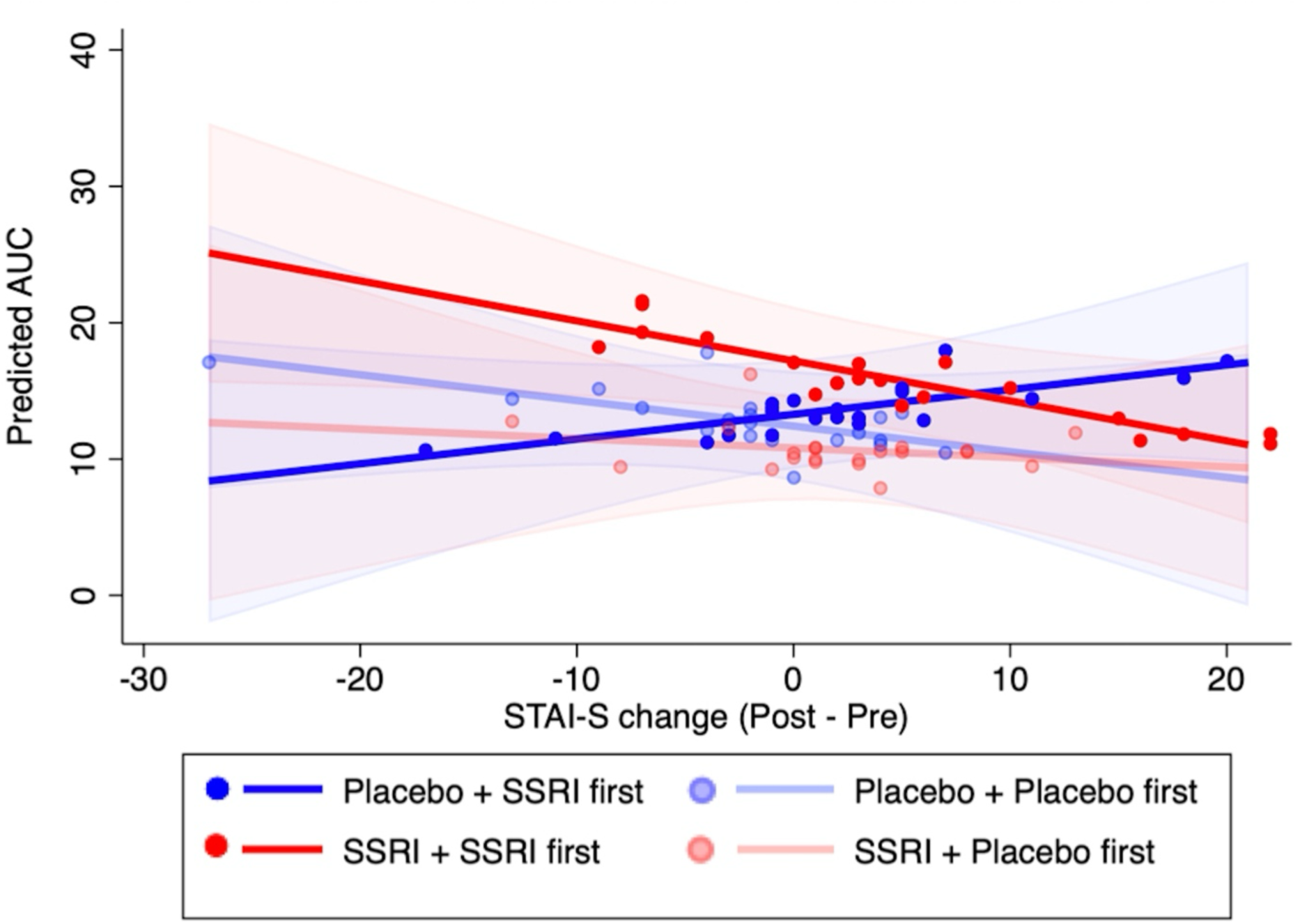
Estimated simple slopes for the relationship between state anxiety change and persistence, plotted separately for each medication condition and condition order, with individual adjusted data points. Shaded areas represent 95% confidence intervals. When participants had taken the SSRI, and it was given at a test visit prior to the placebo test visit, they were willing to wait longer for delayed rewards, but the size of the effect depended on how much participants’ anxiety increased over the course of the session. SSRI = selective serotonin reuptake inhibitor. AUC = area under the survival curve from the persistence task. STAI-S = Spielberger State Anxiety Inventory - State version.

### Exploratory Analyses

#### Tests for potential mechanism

Since we found support for the hypothesis that serotonin increases persistence, we next examined the effects of the SSRI on behavior in foraging (for details, see *Methods* and Fig. 7), time perception, temporal discounting, and risk tolerance tasks, to assess the evidence for different potential mechanisms by which serotonin might increase persistence.

There was no effect of the SSRI on foraging behavior. The dependent variable in the foraging task was the “leaving threshold,” the average number of berries in the last harvest that participants made in a patch before moving on to the next patch. When this value is lower, that indicates that people are staying in the patch longer. Since there were two blocks with two different travel times, we included Travel Time (Long / Short) as a covariate. We also included Block Order (Long first / Short first) as a covariate. The effect of SSRI on leaving threshold failed to reach significance (β = -0.47; 95% CI [-0.97, 0.04]; *p* = 0.070; Supplementary Table 2). Nevertheless, it is noteworthy that the effect of serotonin was in the predicted direction, with participants staying in patches longer after having taken the SSRI (Cohen’s d_z_ = -0.29; 95% CI: [-0.60, 0.02]). This result did not change when we ran a post-hoc analysis that also included ΔSTAI, condition order, and all two-way and three-way interactions between those covariates and medication in the model (β = -0.29; 95% CI [-1.07, 0.50]; *p* = 0.471). Therefore, we did not consider leaving threshold as a potential mediator in our persistence model.

Participants were responsive to opportunity costs in the foraging task, however (Supplementary Table 2). As expected, there was a significant effect of Travel Time on the leaving threshold (β = -1.42; 95% CI [-1.82, -1.02]; *p* < 0.001), with participants leaving patches later when travel times were longer (estimated marginal means for leaving threshold in the long travel time block = 3.14; short travel time block = 4.19). There was also a significant effect of Block Order (β = -0.78; 95% CI [-1.35, -0.21]; *p* < 0.001), which was driven by a Block Order x Travel Time interaction (β = 0.89; 95% CI [0.32, 1.46]; *p* = 0.002). Specifically, when the travel time was long, the block order did not affect the leaving threshold (mean difference between block orders = 0.26; 95% CI [-0.25, 0.78]; *p* = 0.314). When the travel time was short, however, then behavior was influenced by the block order (mean difference between block orders = −0.55; 95% CI [-1.05, -0.05]; *p* = 0.031); people stayed in patches longer if the short travel time block was presented after the long travel time block. Therefore, people’s tendency to stay in a patch longer carried over from the long travel time block to the short travel time block, but their tendency to leave a patch earlier did not carry over from the short travel time block to the long travel time block.

There were no effects of the SSRI on any of the other tasks. There were no effects of the SSRI on time perception accuracy (β = -0.03; 95% CI [-0.12, 0.05]; *p* = 0.405; Cohen’s d_z_ = -0.13; 95% CI: [-0.43, 0.18]), temporal discounting (β = 0.22; 95% CI [-0.16, 0.61]; *p* = 0.260; Cohen’s d_z_ = 0.17; 95% CI: [-0.13, 0.48]), or risk tolerance (β = 0.003; 95% CI [-0.07, 0.07]; *p* = 0.942; Cohen’s d_z_ = 0.01; 95% CI: [-0.29, 0.31]; see *Methods* for full model specifications and details). Therefore, we did not consider any of these variables as potential mediators in our persistence model.

#### Age modulates effects of L-dopa on foraging

For completeness, we performed unplanned exploratory analyses of the effects of L-dopa on behavior in the foraging, time production, temporal discounting, and risk tolerance tasks. L-dopa did not influence time perception accuracy (β = -0.02; 95% CI [-0.11, 0.07]; *p* = 0.682; Cohen’s d_z_ = - 0.06; 95% CI: [-0.37, 0.24]), temporal discounting (β = 0.16; 95% CI [-0.21, 0.53]; *p* = 0.406; Cohen’s d_z_ = 0.13; 95% CI: [-0.18, 0.43]), or risk tolerance (β = 0.003; 95% CI [-0.09, 0.10]; *p* = 0.951; Cohen’s d_z_ = 0.01; 95% CI: [-0.29, 0.31]; see *Methods* for full model specifications and details).

L-dopa also had no effect on leaving threshold in the foraging task (β = -0.30; 95% CI [-0.79, 0.19]; *p* = 0.223; Cohen’s d_z_ = -0.09; 95% CI: [-0.39, 0.22]). There was a medication x travel time interaction in this model (β = 0.45; 95% CI [0.02, 0.87]; *p* = 0.039), however, such that L-dopa reduced the extent to which travel time influenced the leaving threshold (difference in marginal means between block types in leaving threshold on placebo = 0.97; on L-dopa = 0.75; Supplementary Fig. 2).

We then ran these same exploratory analyses again, but this time applying the same covariate selection procedure that we used for the main persistence task analyses. The only positive result was an effect of L-dopa on the leaving threshold in the foraging task that emerged when controlling for age (β = 2.13; 95% CI [0.49, 3.78]; *p* = 0.011; Table 4). In this model, there was a significant medication x age interaction (β = -0.10; 95% CI [-0.17, -0.04]; *p* = 0.003; Fig. 6a). A simple slopes analysis on the relationship between age and leaving threshold for each medication condition (adjusted for block type and foraging order) revealed a significant association between age and leaving threshold when people were on L-dopa (β = -0.06; 95% CI [-0.11, -0.01], *p* = 0.017), but not on placebo (β = 0.03; 95% CI [-0.01, 0.07], *p* = 0.179). Thus, younger people left patches earlier (i.e., explored more) after taking L-dopa, and older people explored less after taking L-dopa.

**Fig. 6.**
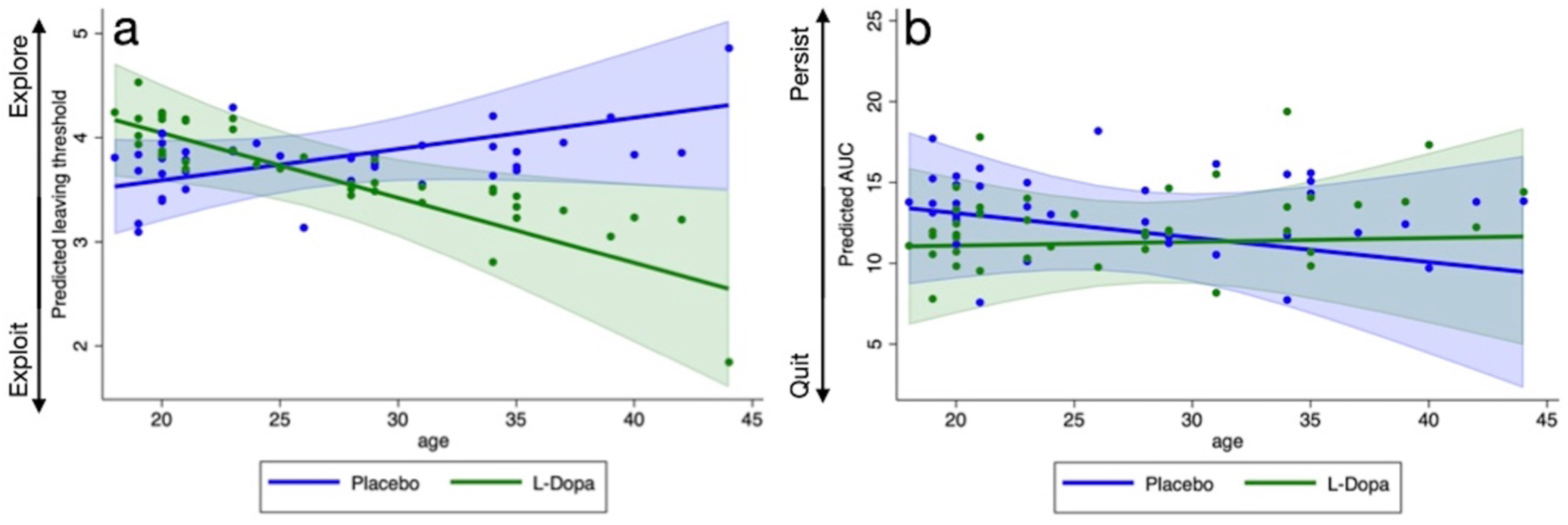
Interactions between age and medication condition on foraging (a) and persistence (b). (A) Estimated simple slopes for the relationship between age and leaving threshold (i.e., number of berries at last harvest), plotted separately for the L-dopa and placebo conditions, with individual adjusted data points. Data are averaged across the long and short travel time blocks. There was a significant interaction between age and medication condition (β = -0.10; 95% CI [-0.17, -0.04]; *p* = 0.003). (B) Estimated simple slopes for the relationship between age and persistence, plotted separately for the L-dopa and placebo conditions, with individual adjusted data points. While the interaction between age and medication condition did not reach significance (β = 0.17; 95% CI [-0.01, 0.36]; *p* = 0.068), there was a significant effect of L-dopa on persistence after controlling for age (β = -5.50; 95% CI [-10.70, -0.30]; *p* = 0.038). AUC = area under the curve. Shaded areas represent 95% confidence intervals.

**Table 4.** Fixed effects of L-dopa on leaving threshold in foraging task without age (Model 1), and with age as an additional predictor (Model 2)

| Predictor | Model 1 |  | Model 2 (with age) |  |
| --- | --- | --- | --- | --- |
| | $\beta$ | 95% CI | $\beta$ | 95% CI |
| L-dopa<br>(1 = L-dopa; 0 = Placebo) | -0.30 | -0.79, 0.19 | 2.13* | 0.49, 3.78 |
| Travel Time<br>(1 = Long; 0 = Short) | -1.42*** | -1.80, -1.04 | -2.33** | -3.84, -0.83 |
| L-dopa x Travel Time | 0.45* | 0.02, 0.87 | 1.33 | -0.29, 2.96 |
| Block Order<br>(1 = Long first; 0 = Short first) | -0.78* | -1.38, -0.18 | -0.97 | -3.55, 1.61 |
| L-dopa x Block Order | 0.20 | -0.49, 0.89 | -1.27 | -3.65, 1.12 |
| Travel Time x Block Order | 0.89** | 0.35, 1.43 | 1.04 | -1.14, 3.23 |
| L-dopa x Travel Time x Block Order | -0.45 | -1.05, 0.16 | 0.54 | -1.81, 2.90 |
| Age |  |  | 0.31 | -0.06, 0.08 |
| L-dopa x Age |  |  | -0.10** | -0.17, -0.04 |
| Travel Time x Age |  |  | 0.04 | -0.02, 0.10 |
| L-dopa x Travel Time x Age |  |  | -0.04 | -0.10, 0.03 |
| Block Order x Age |  |  | 0.01 | -0.09, 0.10 |
| L-dopa x Block Order x Age |  |  | 0.07 | -0.02, 0.16 |
| Travel Time x Block Order x Age |  |  | -0.01 | -0.09, 0.07 |
| L-dopa x Travel Time x Block Order x Age |  |  | -0.03 | -0.11, 0.06 |

Note: Dependent variable is leaving threshold (average number of berries at last harvest) from foraging task, with higher values indicating leaving earlier. In Model 1, random effects were included for the intercept, L-dopa, and Travel Time Block terms (the model did not converge when also including a random effect on Block Order). In Model 2, random effects were included for the intercept, L-dopa, Travel Time Block, Block Order, and age terms. CI = confidence interval. \* $p < 0.05$ , \*\* $p < 0.01$ , \*\*\* $p < 0.001$ .

In light of this result, we ran an exploratory mixed-effects regression examining the effects of L-dopa and age and their interaction on persistence (with random intercepts and random slopes on medication and age, and with no other covariates in the model). In this regression, there was a significant effect of L-dopa on AUC (β = -5.50; 95% CI [-10.70, -0.30]; *p* = 0.038). There was a trend toward a medication x age interaction (β = 0.17; 95% CI [-0.01, 0.36]; *p* = 0.068; Fig. 6b), but no main effect of age on AUC (β = -0.15; 95% CI [-0.56, 0.26]; *p* = 0.469). Therefore, L-dopa reduced persistence after accounting for effects of age. The marginal medication x age interaction appeared to be driven by reduced persistence on L-dopa in younger adults and increased persistence on L-dopa in older adults.

## Discussion

In this pre-registered, placebo-controlled, double-blind, crossover pharmaco-behavioral study, we examined the effects of a single dose of L-dopa (a dopamine precursor drug) and escitalopram (an SSRI) on persistence for delayed rewards in humans. We expected that the dopaminergic medication would reduce persistence (Hyp. 1), and that the serotonergic medication would increase persistence (Hyp. 2). We found no support for the first hypothesis of an effect of L-dopa, but we did observe the expected effect of the SSRI on persistence. This effect was qualified by two interactions: the SSRI increased persistence more when the SSRI was given prior to the placebo, and when participants showed no increase in state anxiety after taking the medication. There were no significant effects of medication on the other tasks we included, which measured patch-foraging, time perception, temporal discounting, and risk tolerance. However, in two exploratory analyses in which we controlled for age, L-dopa increased exploration in the patch-foraging task and reduced persistence in the persistence task. Younger adults showed effects in the expected direction, but these effects tended to go away or even reverse in participants who were relatively older.

Previous theoretical and experimental work^7,22^ has shown that decisions about persisting for delayed rewards structurally resemble explore/exploit decisions in foraging contexts. It has been proposed that, in foraging, dopamine tracks the average reward rate, or the perceived opportunity cost of time. Therefore, our first hypothesis was that L-dopa would reduce persistence in our persistence task. Moreover, we also expected that exploratory analyses of mechanism would reveal increased exploration after L-dopa in a patch-foraging task. In our pre-registered analysis, we found no effect of L-dopa on persistence. Nevertheless, we tested whether L-dopa increased exploration in foraging, and while it had no overall effect, there was a medication x travel time interaction, such that, on L-dopa, the difference in leaving threshold between long and short travel time blocks was attenuated. While this finding goes against the notion that dopamine carries an opportunity cost of time signal^27,29^, it is consistent with other work showing that pharmacological manipulations of dopamine change the extent to which external manipulations of opportunity cost (such as the travel time manipulation here) influence decision making. One study^30^ found that a D2 receptor agonist, cabergoline, reduced the effect of environment reward rate on choice, and another study^31^ found that a D2 receptor antagonist, amisulpride, increased the effect of environment reward rate on choice. Although those studies used different pharmacological agents, our finding adds to a line of research suggesting that dopamine underlies flexible dynamic decision making in response to changes in opportunity cost.

Even more intriguing, however, were our exploratory analyses suggesting that L-dopa reduced persistence and increased exploration in an age-dependent manner. There was a significant interaction between age and L-dopa on foraging, such that younger adults explored more on L-dopa, while older adults explored less. In our persistence task, this interaction did not reach significance, but trended in the same direction. These findings should be interpreted cautiously, as they were exploratory, and our study was not designed to test for age effects. Furthermore, our sample was composed of mostly young adults (age range: 18-44). Nevertheless, ample research has shown that the effects of dopamine on decision making may be nonlinear, and may depend on a person’s baseline dopamine receptor sensitivity^67^. Given that there are complex changes in the dopamine system over the course of adulthood^68,69^, age may be a critical factor to consider in this line of research going forward.

We found support for our second hypothesis: an increase in serotonin transmission increased people’s willingness to persist through experienced delays to obtain rewards. To our knowledge, this is the first demonstration that an SSRI increases persistence in humans. This finding is consistent with a growing body of literature in rodents^24,26,37^ and macaque monkeys^36,41^, however, with some research suggesting that the mechanism by which serotonin influences decision-making is via interactions between the medial prefrontal cortex and the dorsal raphe nucleus^39,70^.

Our result is in line with the theory that serotonin signals the inverse of the average reward rate of the environment, thus leading people to stay at a current resource for longer^32,33^. However, it is also consistent with an alternative theory, which posits that serotonin increases patience by changing the time horizon over which anticipated future outcomes influence current decisions^47,71^. We included a foraging task and a temporal discounting task in order to distinguish between these accounts and probe the mechanism by which the SSRI increases persistence. However, our results were not definitive. Though there was a trend towards serotonin increasing exploitation in foraging, this effect was not statistically significant, and there was no effect of serotonin on temporal discounting. These results might indicate that serotonin’s role in reducing impatience may be secondary to its role in promoting exploitation.

However, there are several other theories of serotonergic function that have been proposed in recent years, which may also be consistent with our findings. Harkin and colleagues^72^ recently proposed that serotonin carries a prospective code for value. This idea is consistent with research showing that serotonin neurons ramp up their firing during delays until rewards are received in trace conditioning^72^. Our results are consistent with this idea, since an increase in the prospective value of the reward for which one is currently waiting would increase willingness to wait for that reward, when opportunity costs are kept constant. Serotonin may also signal the availability of time and resources, such that increased serotonin would promote the willingness to “spend” those resources^73^. In our paradigm, this might manifest as an increased willingness to wait. Within a predictive coding framework, serotonin could track uncertainty in the brain’s predictions, a possibility that is supported by recent work in zebrafish^74^. That the effects of the SSRI were strongest when participants were first gaining experience with the task, when uncertainty was likely highest, could speak to a role for serotonin in updating wait time predictions based on feedback. Finally, another recent theory of serotonergic function suggests that serotonin is crucial for “meta-decision making”^75^; it integrates information about costs and benefits over time in order to change an animal’s overall strategy or motivational state^41^. This process takes into account “meta” factors about the environment, including how controllable and reward-rich it is. Thus, the effects of serotonin on persistence may depend on the controllability and reward-richness of the environment. It will be worthwhile to test how serotonin affects persistence in different contexts that have different reward timing statistics. In general, more research is needed to better understand the precise mechanism by which, and the circumstances under which, serotonin increases persistence for delayed rewards in humans.

The extent to which serotonin increased persistence depended on two factors: condition order and state anxiety. First, those participants who received the SSRI before they received the placebo were more likely to show the effect. This result makes sense in light of the finding that participants tend to wait less time on average the second time they do the persistence task (see Pilot Data in *Methods*). One possible reason for this practice effect is that participants update their waiting policy more from rewarded trials than from quit trials^76^. This asymmetry in updating may bias participants toward waiting less time as they gain experience, since they can observe rewards that occur sooner than their current willingness-to-wait limit, while rewards that would have occurred after their current willingness-to-wait limit are not observed. Since two-thirds of participants received the SSRI either at the second or third visit, this effect of previous experience most likely counteracted the effect of the drug. Although this analysis was underpowered, it is notable that the fourteen participants who were given the SSRI on the first day tended to wait longer (average AUC = 16.8) than those who were given placebo on the first day (average AUC = 12.7). Second, those participants who showed a greater increase in state anxiety after taking the SSRI were less likely to show increased persistence. More research is needed to understand the precise role that anxiety plays in persistence, since anxiety was unrelated to decision making in the placebo condition. It is possible that the bodily symptoms evoked by the SSRI (e.g., dizziness, nausea) caused some participants to both feel more anxious after taking the drug and to be less willing to wait through delays while experiencing those symptoms.

This study has a few limitations. First, although the study was sufficiently powered to detect meaningful effects, our sample size was relatively small (*n* = 42), so it is possible that some null results are due to a lack of power. Indeed, although we cannot conclude that L-dopa reduced persistence, we also cannot conclude that behavior in the L-dopa and placebo conditions was statistically equivalent. Second, we did not perform blood tests to ensure that the medications increased plasma concentrations of serotonin or dopamine; however, many previous studies using these doses or smaller doses have shown that they are sufficient to influence behavior. Third, there is compelling research to suggest that dopamine can increase the willingness to bear effort costs^77^, and that the serotonergic system may be responsive to effort costs as well^75^, but we did not include an effort-based decision-making task to explore this potential mechanism. Fourth, our persistence task did not manipulate the opportunity cost of time, so we could not test if dopamine or serotonin would have changed the extent to which people responded to such manipulations. Finally, using a within-subjects design meant that participants performed the same tasks three times. Thus, effects of previous experience may have interfered with our ability to detect medication effects.

These limitations are outweighed by several notable strengths. First, we pre-registered our hypotheses, sample size, inclusion and exclusion criteria, and entire analysis plan. Therefore, we can be confident that our conclusions do not depend on “researcher degrees of freedom.” Second, using a within-subjects design allowed us to avoid selection bias, as well as to demonstrate that a person’s persistence for delayed rewards can be manipulated through a change in serotonin levels. Finally, our task was designed so that earnings were equivalent regardless of how long people persisted. Therefore, changes in persistence from one session to the next cannot be attributed to changes in motivation or learning.

In sum, here we showed that a single 20 mg dose of escitalopram, an SSRI, increases persistence for delayed rewards. This is a novel finding in humans that adds to a growing literature on links between serotonin and behavioral persistence in non-human animals. We also provided tentative evidence that L-dopa may influence dynamic decision making in an age-dependent manner. Future research should consider how chronic use of serotonergic and dopaminergic drugs affects these processes, how these processes are impacted by psychopathology, and what the implications of these effects are for real-world decisions.

## Methods

### Ethics information

This research complies with all relevant ethical regulations and was approved by the Institutional Review Board of the University of Pennsylvania (IRB #843535). Informed consent was obtained from all participants. Participants were compensated at least ∼$265 if they completed the full protocol. Participant payments were prorated as follows: $15 for an initial intake visit, $40 for each of three test visits, and a $100 completion bonus. In addition, participants received bonus payments for performance on the persistence task and foraging task (described below). These bonus payments totaled ∼$15 per person (∼$5/visit) in the persistence task and ∼$15 per person (∼$5/visit) in the foraging task. Participants also had the opportunity to receive a bonus based on their choices in the Monetary Choice task, but the receipt of this bonus and its value was probabilistic (described below). All payments were delivered using the Greenphire ClinCard (debit card) system.

### Design

The study used a within-subjects placebo-controlled double-blind design. Procedures took place in the Outpatient Study Unit of the Center for Human Phenomic Science at the Penn Presbyterian Medical Center. The study involved four visits: an intake screening visit and three test visits, separated by at least 7 days. See Figure 1 for schematic of the procedure.

#### Intake Visit

The primary purpose of the intake visit was to assess if participants were eligible for the study, according to our inclusion and exclusion criteria (see *Sampling Plan*). After informed consent was obtained for all study visits and procedures, a medical assessment was performed by a nurse practitioner. This assessment included a detailed medical (including psychiatric) history and physical exam including height, weight, and vital signs (pulse, blood pressure, temperature, and respiration rate). In addition, female participants took a pregnancy test, and all participants completed a standard urine drug screen to assess for levels of cocaine, marijuana, opiates, amphetamine, methamphetamine, barbiturates, benzodiazepines and phencyclidine. If participants tested positive for any drug other than marijuana, they were excluded from enrolling in the study. If they tested positive for marijuana, they were asked about their last marijuana use, and they were included in the study if their last marijuana use was > 24 hr before the session. People with a history of substance abuse or dependence within the last 6 months were excluded from the study. All participants were asked to refrain from alcohol, nicotine, and marijuana use for 24 hours before each test visit. In some cases, if requested by the participant and compatible with scheduling, the Intake procedures were performed on the same day as the first test visit. Otherwise, the schedule of test visits was determined at the end of the intake assessment, if the participant fulfilled inclusion criteria. An attempt was made to schedule test visits at the same time each day, and task administration times were recorded, in order to control for circadian effects on task performance and/or medication absorption. During scheduling, participants were asked not to eat a large meal for ∼2 hours prior to each test visit, in order to minimize differences in drug absorption between subjects.

#### Test Visits

##### General procedure, randomization, and blinding

The procedures described below were repeated at all three test visits, with the exception that the medications or placebos administered (L-dopa, escitalopram, or placebo) were different on the three days. There were six possible orders across the three test visits: (1) L-dopa-Escitalopram-Placebo, (2) L-dopa-Placebo-Escitalopram, (3) Escitalopram-Placebo-L-dopa, (4) Escitalopram-L-dopa-Placebo, (5) Placebo-L-dopa-Escitalopram, and (6) Placebo-Escitalopram-L-dopa. The order was counterbalanced across subjects before data collection began. Experimenters and participants were blind to the order in which the medications and placebos were administered.

The procedures were as follows. When participants arrived for their test session, their vital signs were taken, and female participants took a pregnancy test. Participants also reported on any changes in their medication or drug use since the previous visit. Participants then filled out the Positive and Negative Affect schedule (PANAS^78^) and the Spielberger State Anxiety Inventory (STAI-S^79^) to measure their affective state. Analyses with state questionnaire data were exploratory, with the exception that if there were significant changes in affect following administration of L-dopa or escitalopram, the change in affect was included as a covariate in analyses (see *Control Analyses* and Supplementary Table 1). Immediately following this, participants took the first medication or placebo. Approximately every hour after that, up until discharge 4 hours later, a nurse assessed and recorded their vital signs (heart rate, respiration rate, temperature, and blood pressure) to monitor participants for safety. Ninety minutes after the first drug or placebo was administered, participants took a second drug or placebo, and were given a standardized snack (e.g., a granola bar). Finally, ninety minutes after that (3 hr after the initial drug or placebo was administered), participants performed the experimental tasks (described below). After the tasks were finished, they once again completed the PANAS and STAI-S questionnaires and they were asked to guess which condition they were in (to assess blinding). Their vital signs were assessed, and they were also asked to report any side effects or sensations before being discharged, when discharge was deemed safe. In addition, at the end of the last test visit, participants were asked to guess the order in which they were given the two medications and placebo. At the end of the last test visit, they were also asked about strategy use and if they suspected that the experimenter wanted them to behave in a certain way.

##### Medication administration procedure

All medications were provided, stored, prepared (randomized, blinded), recorded, dispensed, and re-collected/destroyed by the Investigational Drug Service (IDS) at the University of Pennsylvania. IDS is the research pharmacy serving the University of Pennsylvania community, and it is involved in almost every clinical trial. With their state-of-the-art inventory management system, IDS ensures that all investigational drugs are procured, stored, inventoried, dispensed and used in compliance with all relevant federal and local regulations, and in accordance with Hospital, Medical Center and University policies and guidelines. For this study, all medication orders were documented in a centralized electronic medical records system, and all capsules were packaged with a unique barcode that linked them to information about the study visit.

Because the pharmacokinetics of escitalopram differ from L-dopa, we used a multiple-placebo procedure to ensure blinding. Escitalopram reaches peak levels ∼3 hours after it is administered, whereas controlled-release L-dopa (the formulation we used) reaches peak levels ∼1.5 hours after it is administered. Moreover, the elimination half-life of levodopa is only about 1.9 hours^80,81^ (compared to 27-33 hours for escitalopram)^82,83^, so its concentration would not stay near peak levels for long enough to warrant giving it at the same time as escitalopram. Therefore, we designed our study such that participants did the experimental tasks 3 hr after taking escitalopram (at the escitalopram visit), and 1.5 hr after taking L-dopa (at the L-dopa visit). Timing the procedure in this way allowed the drug concentrations to be near-peak for the entire experimental task battery, which took about 40 minutes to complete.

At the beginning of each test visit, subjects received either a single oral dose of 20 mg of the SSRI escitalopram (if it was the escitalopram visit) or placebo otherwise. Based on previous studies, we intended to use 30 mg of citalopram^28,84,85^, which is a high enough dose to ensure that serotonin transmission is increased^86^, but low enough that side effects are minimal. Our university pharmacy did not have 30 mg of citalopram available in single tablets, however; thus, we opted for 20 mg of escitalopram. Escitalopram is the S-enantiomer of citalopram and is more potent, so 20 mg of escitalopram is about equal to 40 mg of citalopram. It has also been used at this dosage both in the clinic (trade name: Lexapro) and for psychological research^42^.

Two hours later, if it was the L-dopa visit, participants received a single oral dose of controlled-release 200 mg levodopa + 50 mg carbidopa (trade name: Sinemet CR). Otherwise, they received a placebo. Carbidopa is often paired with levodopa in clinical settings. It serves to prevent the breakdown of levodopa in the bloodstream, resulting in more levodopa entering the brain. (Note that throughout the manuscript, we refer to this combination of levodopa and carbidopa as “L-dopa.”) Based on previous studies of dopamine and decision-making^28,65,87^, we intended to use 150 mg levodopa + 37.5 mg benserazide (which is, like carbidopa, a peripheral inhibitor of the DOPA decarboxylase enzyme that metabolizes levodopa). This formulation was not available at our university pharmacy, so we opted instead for the comparable dose of 200 mg levodopa + 50 mg carbidopa. While this dose is slightly higher than that used in other studies, it does not affect mood^88^. We also opted for the slightly higher dose of levodopa to more closely match the potency of the serotonergic medication.

We decided not to include blood tests to assess plasma levels of these medications. First, previous research has shown that the doses that we are using here will result in significantly elevated plasma levels of escitalopram and L-dopa at the expected time points in healthy young subjects^81–83^. Second, blood tests are not deemed necessary when the risk of participant non-compliance is very low^86^, as it is here. Indeed, previous similar studies using these compounds did not include blood tests^44,65^. Third, blood tests are costly, both from a financial perspective (requiring additional nursing services, equipment, and laboratory services) and from the participant’s perspective, since they are often perceived as aversive. Finally, even if we did determine plasma levels of the drug, that does not guarantee that the levels of the drug that are psychoactive would be sufficient to affect behavior^89^. Plasma levels of escitalopram and L-dopa are often weakly or not correlated with behavioral effect sizes^42,90^, even when there are average effects of drug on behavior. We did take some measures to protect against the possibility that the doses would not be sufficient, however; we selected doses that are somewhat higher than the average doses used in the literature, and we made plans (see *Analyses* below) to examine effects of body mass index and sex, which might impact the extent to which the medications are psychoactive.

We carefully considered other possible behavioral or physiological measures that could serve as a positive control. Unfortunately, neither of these pharmacological compounds has a significant impact on vital signs^81,83^, so we did not have the option of using a less invasive correlate of drug levels, such as heart rate or blood pressure. Moreover, to our knowledge, the effects of L-dopa and escitalopram on existing behavioral tasks are not well enough established such that we could use another task as a positive control.

To summarize, at each test visit, participants received either two placebo pills (if it was their placebo visit) or one active pill and one placebo pill. Escitalopram, L-dopa, and placebo capsules were packaged to be indistinguishable, to maintain double-blinding.

##### Baseline Assessments of Personality and Cognitive Ability

At the first test visit, while participants were waiting after receiving the first dose of medication (escitalopram or placebo), they completed a set of baseline assessments of subclinical psychopathology, personality, and cognitive abilities.

One set of questionnaires assessed subclinical levels of psychopathology, including the degree to which participants experience anhedonia, depression, anxiety, attention deficit hyperactivity disorder (ADHD) symptoms, and compulsivity. These questionnaires were included for the purposes of future research, particularly given our interest in the relationships between depression, anxiety, and decision making^91,92^. These questionnaires included: the Motivation and Pleasure Scale – Self-Report scale^93^, the Quality of Life Enjoyment and Satisfaction Questionnaire^94^, the Beck Depression Inventory^95^, the Beck Anxiety Inventory^96^, the Depression Anxiety and Stress Scale^97^, the Spielberger Trait Anxiety Inventory^79^, the Adult ADHD Self-Report Scale^98^, the UPPS-P Impulsivity questionnaire^99^, and the Obsessive-Compulsive Inventory^100^.

The Barratt Impulsiveness Scale^101^ (BIS-11) assessed trait impulsivity. This questionnaire was included given previous research showing that baseline levels of impulsivity, as measured by the BIS, can modulate the effects of L-dopa on decision-making^53^. If impulsivity were to influence our primary outcome variable, either on its own or in interaction with L-dopa, then we planned to include it as a covariate in the primary analysis.

In the same vein, we included the automated operation span working memory task^102^, a standard measure of individual differences in working memory capacity, since previous studies have shown that the effects of dopaminergic medications on behavior can depend on baseline working memory capacity^103^. Working memory capacity was also considered as a potential covariate.

Finally, participants completed the matrix reasoning and similarities subtests of the Wechsler Abbreviated Scale of Intelligence (WASI-II^104^) as a measure of general cognitive ability. These tests were included for future research purposes.

##### Experimental tasks

At the 3 hr mark of each test visit, participants completed a persistence task. Then they completed a foraging task, a time production task, and a monetary choice task (which included both intertemporal choice and risky choice questions). While the persistence task was always completed first, the remaining three tasks were given in a random order, which was counterbalanced across subjects (6 different orders). Each participant saw the same order of tasks at each visit, however. Each task took approximately ten minutes to complete.

*Persistence task.* Participants performed a 10-minute block of a validated persistence task at each test visit (Fig. 2). This task structure has been previously used to study willingness-to-wait^18,22,105^ and was programmed using the Psychophysics Toolbox in Matlab^106,107^. Participants’ goal in this task was to earn as much money as they could in 10 min, since they kept any money that they earned. On each trial, participants saw a token labelled “0¢” on the screen. After a variable delay (up to 40 s), the token “matured” and was then worth 10¢. At that point, participants could press the spacebar to sell the token (get 10¢), and they accumulated earnings throughout the task. Participants could also sell the token *before it matured* if they felt that it was taking too long and they wanted to move on to a new token, even though they would not earn any money from selling 0¢ tokens. When they sold the token, the word “SOLD” appeared over the token and the task advanced to the next trial, with a new 0¢ token and updated earnings. The value for which they sold the token was added to the participant’s total earnings. Total earnings were displayed on the screen throughout the experiment. Participants did the same persistence task, with the same timing parameters, at every test visit. However, the color of the token pre-maturation was different each time (green at Test Visit 1, purple at Test Visit 2, and magenta at Test Visit 3). The color change was meant to serve as a subtle cue that the timing distribution may have changed since the previous visit. However, the color was never pointed out to participants, and participants were never told whether or not there was a change in the timing distribution.

The delays until the token matured were drawn from a distribution that yields an approximately equivalent reward rate regardless of the participant’s waiting policy (Supplementary Fig. 1). The waiting policy is defined as the time at which a decision maker will give up waiting on each trial if the reward has not yet arrived. The expected return for giving up at time *t* is calculated as follows. Let *p*_t_ be the proportion of rewards delivered earlier than *t*. Let τ_t_ be the mean duration of these rewarded trials. One trial’s expected return, in dollars per second, is:

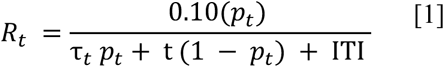

The numerator is the trial’s expected gain in dollars, given a 10¢ reward. The denominator is the trial’s expected time cost in seconds, which depends on the length of the inter-trial interval (ITI). As the ITI increases, waiting policies that are very short will yield lower expected reward rates. Here we used a 0.25 s ITI, which allowed for participants to get feedback when selling the token, but was sufficiently short that very short waiting policies were not severely disadvantaged.

Under perfect conditions – that is, with no intertrial interval, and with an infinite range of possible reward times – a “flat” expected return rate across all waiting policies could be obtained with an exponential reward timing distribution. Here instead we used a more flexible Generalized Pareto distribution in order to approximate a flat relationship between waiting policy and expected return rate, given our experimental conditions. The Generalized Pareto distribution has the following cumulative distribution function:

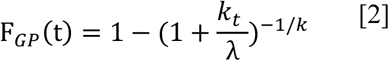

(Here we omit the location parameter θ, which we set to zero, implying that zero is the shortest delay.) The parameter *k* is the shape parameter. When *k* = 0 (and θ = 0), the generalized Pareto distribution equals the exponential distribution with scale parameter λ. Increasing *k* slightly allowed us to optimize our design further. We simulated R_t_ (Eq. [1]) for waiting policies *t* in the range [0, 40] s in increments of 0.01 by taking 100,000 draws from a Generalized Pareto distribution. We did this for each *k* in the range [0.001, 0.1] in increments of 0.001. We then found the value of *k* that resulted in the minimum absolute slope of a linear fit between *t* and R_t_ over the range of 5 s to 40 s. Therefore, the final reward timing distribution was a Generalized Pareto distribution with the following parameters: *k* = 0.034; λ = 12; θ = 0. To avoid very long delays, the distribution was truncated at 40 s, such that any delays longer than 40 s were set to 40 s. Furthermore, as in previous studies^22,105^, to ensure that participants’ experiences would be representative of the true underlying distribution, delays were not drawn randomly on each trial, but instead were sampled from each octile of the distribution in random order before an octile was repeated.

To quantify participants’ persistence, we constructed a Kaplan–Meier survival curve for each participant, which plots, for each time *t* into the waiting interval, the participant’s probability of waiting at least until *t* if the reward is not delivered earlier. The area under this survival curve (AUC) represents the average number of seconds an individual was willing to persist. We computed the AUC for each participant for each session.

#### Pilot data

Since participants performed this task three times, we first established that behavior was stable over time in the absence of any manipulations. In a pilot experiment, participants (*n* = 32; mean age = 26.07; SD = 7.30; range: 18-56; 17 F, 14 M, 1 not reported) performed a 10-min block of the persistence task twice, spaced one week apart. Sampling from the distribution was as described above (i.e., sampled from each octile in random order). The particular delays experienced differed across subjects, and the delays experienced on the first day were not the same as those experienced on the second day. Thirty-one participants completed both sessions and were included in analyses. Test-retest reliability for AUC was moderate in terms of absolute agreement (ICC (3,1) = 0.64; 95% CI [0.31, 0.82]; *p* < 0.001) and high in terms of rank-order consistency (Spearman ρ = 0.85; *p* < 0.001; Supplementary Fig. 3). We also validated that earnings were not correlated with AUC (Day 1 earnings with Day 1 AUC: Spearman ρ = 0.13; *p* = 0.478).

*Foraging task*. In this game^13^, the participant’s goal was to collect as many berries as they could in an 8-min period, which was divided into two blocks lasting 4 minutes each (Fig. 7a). Participants were paid according to how many berries they collected. Specifically, they received in cents the total number of berries divided by 200 (usually ∼$4-$5). On each trial, the participants first hovered their mouse over a center square until it turned green. Then they chose to either “stay” or “leave,” by moving their mouse toward the patch that was directly adjacent to the center box (“stay”) or by moving their mouse to an empty box on the opposite side of the center box (“leave”). Upon choosing to stay, they received a number of berries from the patch as a reward, after a static “handling time” (1 s). The first time they harvested from the patch they received 7 + N (0, 0.25) berries, where N (0, 0.25) is a normal distribution with a mean of 0 and a variance of 0.25. Each time they returned to the same patch, the patch became depleted, and the berry harvest rate declined by 0.5 + N (0, 0.25). On any given trial, the participant could choose instead to “leave.” This brought them to a new, full berry patch. Moving to a new patch took time (“travel time”), thus creating an opportunity cost. In one 4-min block, the patches were spaced close together (travel time was relatively short, T = 3s), and in the other 4-min block (order counterbalanced), the patches were spaced farther apart (travel time was relatively long, T = 10 s). The dependent variable of interest was the patch leaving time, or the number of harvests before the participant left the patch. When this value was larger, that meant that the participant stayed in the patch longer, and *vice versa*. This design is similar to some previous foraging studies, in which the reward received at each harvest depends on the previous harvest’s reward and some noise^10,11^, and the decay rate is the same for each patch (Fig. 7b). Therefore, differences in average reward rate (and thus, opportunity cost) between blocks are driven only by the differences in travel time. This differs from other previous studies in which the decay rate was variable from one block to the next^30^.

**Fig. 7.**
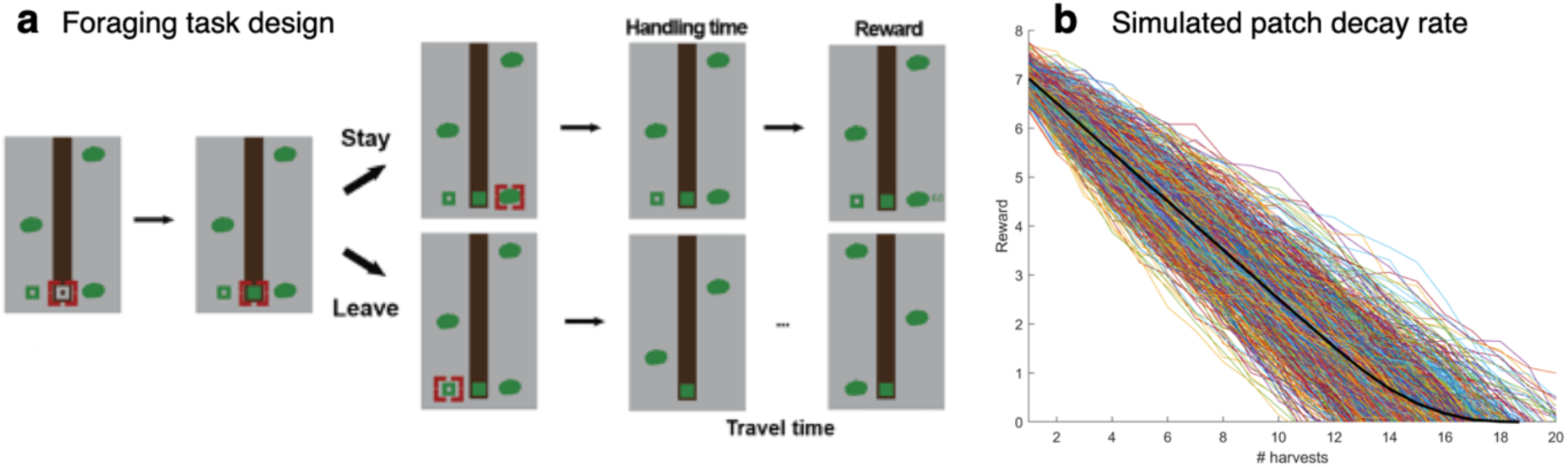
Foraging task. (A) Sample trial. The participant begins each trial by hovering their mouse over the center square until it turns green. Then, they can choose to “stay” in the adjacent patch by moving the mouse in its direction. After a handling time of 1 s, they receive some number of berries (reward). Each successive decision to “stay” results in fewer berries. On any given trial, the participant can choose to leave by moving the mouse away from the patch. This will result in a travel time until the next patch is reached. In one 4-min block, the patches are spaced close together (travel time is relatively short, 3 s), and in the other 4-min block (order counterbalanced), the patches are spaced farther apart (travel time is relatively long, 10 s). (B) Patch decay rate based on 1000 simulations (black line is the average). The number of berries at first harvest is 7 + N (0, 0.25). After this, the number of berries declines by 0.5 + N (0, 0.25) with each harvest.

Behavior in this task was stable over time as well. In a previous experiment, 161 participants performed the foraging task twice, spaced by one day. The reward decrement in the patches encountered by the subjects was not the same in these two rounds of gameplay. However, we observed a high correlation in the average patch leaving time (Spearman ρ = 0.76; *p* < 0.001).

*Time production task.* Each trial of this task^64^ started with the instruction, “Hold down the space bar for XX seconds” (where the amount XX varied across trials). When the participant depressed the space bar, the screen turned blue, indicating that the timer was active. Two practice intervals, for 4 and 8 sec, were performed with feedback, to familiarize the participant with the required movements. Participants completed three trials each for 1, 3, 6, 12, and 24-sec. These intervals are similar to those used in previous studies, and were chosen in order to approximately cover the range of delays that participants experience in the persistence task. Participants were explicitly instructed to resist using counting methods, including rhythmic strategies^108^. At the end of the timing task, participants were asked whether or not they were able to resist using a counting strategy. The outcome of interest was the accuracy coefficient, “theta,” which is the ratio of the estimated duration to the duration requested. Theta values greater than 1 indicate overestimation and lower than 1 indicate underestimation of time intervals.

*Monetary choice task.* Participants completed a monetary choice task with two parts (presented in random order): a risk tolerance block and a temporal discounting block. In the risk tolerance block, participants made a series of 60 choices between a smaller amount of money ($1-$68) available for certain, and a larger amount of money ($10-$100) available with a 50% chance. In the temporal discounting block, participants made a series of 51 choices between a smaller amount of money ($10-$34) to be received immediately, and a larger amount of money ($25, $30, or $35) to be received after a delay (ranging from 1 day to 180 days). This task was self-paced and took ∼10 minutes. This task has been used in several previous studies in our lab, and the question sets reliably capture individual differences in temporal discounting and risk tolerance, with excellent model fit^109–112^.

In order to provide an incentive to participants to choose the options they preferred, while still matching (approximately) the magnitude of possible payments across tasks, we enacted a lottery for the bonus payment from this task. Participants were told that, at each visit, there was a 1/6 chance that we would randomly select one trial from one of the two blocks (risk tolerance or temporal discounting) to be paid out for real. After all of the experimental tasks were completed, participants rolled a standard die to determine if they would get a bonus payment or not. Then, if they were to receive a bonus, the experimenter would flip a coin to determine if the bonus would come from the risk tolerance block or the temporal discounting block. Finally, a random number generator was used to select the random trial to be paid out. If the bonus was from the risk tolerance block and the participant chose the certain amount on the randomly-selected trial, they received that amount that day. If they chose the risky option, the experimenter flipped a coin to determine if they would receive the risky amount or $0. If the bonus was from the temporal discounting block and the participant chose the immediate amount on the randomly-selected trial, they received that amount that day. If they chose the delayed option, then that amount was deposited onto their ClinCard debit card after the delay had passed. Note that since all compensation was received on the debit card, this equates transaction costs for all potential payments.

Each participant’s choice data for the temporal discounting and risk tolerance blocks was fit with the following logistic function using maximum likelihood estimation:

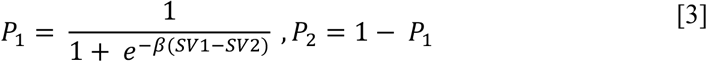

where P_1_ refers to the probability that the participant chose option 1 and P_2_ refers to the probability that the participant chose option 2. SV_1_ and SV_2_ refer to the participant’s estimated subjective value of option 1 and option 2 respectively. The scaling factor β was fitted separately for each task.

In the risk tolerance task, P_1_ is the probability of choosing the risky option, and the subjective values of the options were estimated using a power utility function:

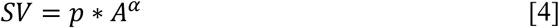

Here *A* is the amount that could be received, *p* is the probability of receipt (*p* = .5 for the risky option, *p* = 1 for the certain option), and α is a risk tolerance parameter that varies across subjects. Higher α indicates greater risk tolerance (less risk aversion).

In the temporal discounting task, P_1_ is the probability of choosing the delayed option, and the subjective values of the options were estimated using a hyperbolic discounting function^113,114^:

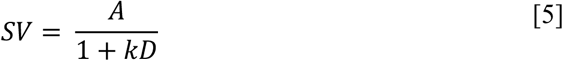

Here *A* is the amount received, *D* is the delay until receipt (for immediate rewards, *D* = 0), and *k* is a discount rate parameter that varies across subjects. Higher *k* indicates higher discounting (less tolerance of delay). The values *k* and α were natural log-transformed before conducting statistical analyses, since they are not normally distributed.

### Sampling plan

#### Inclusion and exclusion criteria

Participants were adults, aged 18-45, in good health as determined on the basis of medical and psychiatric history and examination at the intake visit. In order to qualify, participants could not have had any history of neurological or psychiatric disorders, and they could not have been taking any psychoactive medications. Participants were also excluded if they had done this persistence task in our lab previously (before recruitment, a list of previous subjects were cross-checked to avoid this possibility). They were also required to refrain from using nicotine, alcohol, or illicit drugs (including marijuana) for 24 hours prior to test days, during test days, and for 24 hours after test days. Current intoxication and intoxication within the last 24 hours were assessed via self-report and a urine drug screen at the intake visit, and via self-report only at each test visit. If participants tested positive for any illicit substance other than marijuana at the intake visit, they were not enrolled in the study. If participants reported using nicotine, alcohol, or marijuana during the 24 hours before a test visit, that test visit was rescheduled. The following additional exclusion criteria were also applied:

- Pregnancy (as determined by reported history and reported use of an adequate birth control method as well as urine test on each study day).
- Women who are lactating/breast-feeding.
- History of any clinically significant neurological, psychiatric, metabolic, hepatic, renal, hematologic, respiratory, cardiovascular, gastrointestinal, urological or other medical disorders that might increase risks of L-dopa or escitalopram as determined by the study physician. Active medical conditions that are minor or well-controlled were not exclusionary if they did not affect risk to the patient or metabolism of L-dopa or escitalopram.
- Known hypersensitivity, adverse reaction or contraindication to escitalopram or other SSRIs, or to levodopa/carbidopa, as judged by the study physician.
- Consumption of any grapefruit or grapefruit-containing products within 48 hours before test days (may affect escitalopram metabolism). If participants reported consuming grapefruit or grapefruit-containing products during the 48 hours before a test visit, that test visit was rescheduled.
- Current use or use within 2 weeks of test days of any medications or substances known to or judged by the study physician to cause central nervous system effects interfering with study results, or alter the metabolism or increase the risk of L-dopa or escitalopram, including but not limited to: monoamine oxidase inhibitors (MAOIs), benzodiazepines, barbiturates, other sedatives, antibiotics, sulfonamides, CYP3A4 inhibitors.
- History of active abuse or dependence within the past six months of nicotine, alcohol, or illicit drugs, by DSM-diagnostic criteria and numerical criteria (>=10 cigarettes per day; alcohol > 2 standard units per day [a standard unit equals 12 oz. of beer, 1.5 oz. of 80-proof alcohol or 6 oz. of wine]).

Participants were excluded from analysis post-enrollment if it became clear at any point that they did not fulfill the criteria stated above. One participant was excluded after their second test visit because they were prescribed antidepressants. We also planned to exclude participants if they experienced medical side effects of the medication that reached the level of an “adverse event.” That is, minor side effects that were not too bothersome to the participant were not exclusionary, but we planned to exclude subjects who experienced any side effect that required them to be evaluated by a medical professional before leaving the premises, or that required follow-up by the research team (according to our IRB protocol). Although a few participants experienced nausea and vomiting, these side effects were considered minor and did not qualify as adverse events. Therefore, no participants were excluded on those grounds.

#### Power analysis

The existing research on the effects of L-dopa and escitalopram on foraging-like decisions in humans is limited, so it was not clear what effect size to expect. One recent study, however, used a within-subjects design (as we did here) to test for the effects of a dopaminergic medication on foraging decisions^30^. This study found, using a mixed-effects model, that dopamine affected the use of background reward information to guide patch leaving with a Cohen’s *d*_z_ = 0.425 (a medium effect size), so we powered our study to detect this effect size. We conducted our power analysis using software written by Westfall and colleagues for mixed-effects models^115^, assuming the default settings for standardized variance partitioning coefficients. Given those assumptions, about 45 participants were needed to obtain 95% power to detect an effect size of 0.425. To allow for even numbers in all condition orders, however, and considering financial and time constraints on data collection, we decided to decrease this number slightly to 42 participants. That is, data collection stopped when we had collected full data (three test visits) from 42 participants. According to a power sensitivity analysis, collecting data from this number of participants gave us 94% power to detect an effect size Cohen’s *d_z_* of 0.425, and it gave us >80% power to detect effects at least as large as Cohen’s *d_z_* = 0.339 (small-to-medium).

### Analysis Plan

#### Primary analyses

Hypotheses 1 and 2 were tested using two linear mixed-effects regression analyses in STATA. Model fitting was performed using restricted maximum likelihood, and degrees of freedom were computed using the repeated-measures ANOVA method. The regression specification was as follows, which allowed random intercepts and coefficients on the medication condition factor to vary by subject:

AUC ∼ 1 + medication + (1 | subject) + (1 | subject: medication)

The covariance between random intercepts and coefficients was specified as unstructured. Here *AUC* is the area under the Kaplan-Meier survival curve, our summary statistic of persistence. For the test of Hypothesis 1, only the data from the placebo and L-dopa conditions were analyzed, and *medication* was a categorical predictor with two levels: Placebo and L-dopa. For the test of Hypothesis 2, only the escitalopram and placebo data were analyzed, and the medication condition had two levels: Placebo and Escitalopram. The significance level was set at α = 0.025, to Bonferroni-correct for multiple comparisons. Therefore, we considered Hypothesis 1 to be supported if the coefficient on medication condition was significantly negative (relative to placebo), with *p* < 0.025, and we considered Hypothesis 2 to be supported if the coefficient on medication condition was significantly positive (relative to placebo) with *p* < 0.025.

If the *p*-value for the effect of medication in either of the mixed-effects models was greater than our α level of 0.025, we planned to compare the average AUC between the two conditions of interest (i.e., either L-dopa vs. Placebo or Escitalopram vs. Placebo) using a two one-sided *t*-test equivalence test (TOST^116,117^). This test examines whether behavior in the two conditions is statistically equivalent given a smallest effect size of interest, which we specified as Cohen’s d_z_ = 0.14. This is considered a benchmark value for a “small” effect size in tests comparing dependent means^116^, and in our task would correspond to about a half-second difference in AUC between conditions (we assumed that the data would have similar between-subject variance to our pilot data). If we were to find that both of the two one-sided *t*-tests were significant at *p*<0.05, then we would be able to conclude that behavior between the two conditions is not only not statistically different, but also statistically equivalent.

#### Covariates and control analyses

The inclusion of the following covariates in our primary analysis was contingent on their relationship with the primary variables in the study (see Supplementary Table 1 for details of contingency plan). For the test of Hypothesis 1 with covariates, we conducted the following mixed-effects regression on the subset of data from the L-dopa and placebo conditions (thus, here *medication* was a categorical factor variable with two levels: Placebo and L-dopa):

AUC ∼ 1 + medication * covariate + (1 | subject) + (1 | subject: medication) + (1 | subject: medication: covariate).

Note that here “medication * covariate” encompasses the main effects of medication and the potential covariate, as well as the interaction between them. This same procedure was applied for Hypothesis 2, on the subset of data from the escitalopram and placebo conditions. In the case that either of these more complex models was unable to converge with random effects for the interaction term, we planned to drop random effects terms step-wise until the model did converge. The potential covariates that might be included in our hypothesis tests fell into three categories: (1) variables that may influence persistence or the effects of medication on persistence, (2) variables related to the effectiveness of blinding, and (3) variables related to the participant’s affective state. These are discussed in turn below.

##### Variables that may influence persistence or the effects of medication on persistence

For the Hypothesis 1 analysis (L-dopa vs. Placebo), we entered each of these covariates into this regression in turn: (1) body mass index (continuous variable), (2) age (continuous), (3) sex (binary variable: Female / Male), (4) condition order (binary variable: Placebo condition first / L-dopa condition first), (5) working memory capacity (continuous variable), and (6) Barratt Impulsiveness Scale score (BIS score; continuous variable). If there was a significant main effect of any of these covariates on AUC, or a significant interaction between any of these covariates and medication on AUC (i.e., if the coefficient on the medication x covariate term was significant in the model above at α = 0.05), then we planned to include the relevant covariate(s) in the primary test of Hypothesis 1. In the end, each of these regressions included random intercepts and random slopes for L-dopa only, with three exceptions: the age, sex, and BIS score models additionally included random slopes on age, sex, and BIS score, respectively.

For the Hypothesis 2 analysis (Escitalopram vs. Placebo), here the covariates that were tested serially were: (1) body mass index (continuous variable), (2) age (continuous), (3) sex (binary variable: Female / Male), and (4) condition order (binary variable: Placebo condition first / Escitalopram condition first). Note that we did not expect to see that escitalopram’s effects would be moderated by working memory capacity or trait impulsivity. If there was a significant main effect of any of these covariates on AUC, or a significant interaction between any of these covariates and medication on AUC (i.e., if the coefficient on the medication x covariate term is significant at α = 0.05), then we planned to include the relevant covariate(s) in our primary test of Hypothesis 2. In the end, each of these regressions included random intercepts and random slopes for SSRI only.

##### Effectiveness of blinding

We also wanted to ensure that our blinding was effective and that participants did not know the order that they were given the medications and placebo. If they did know which medication they had taken at the visit, that might influence their behavior in the tasks. At the end of the third test visit, participants were asked to guess the order in which they were given L-dopa, escitalopram, and placebo. Then, the proportion of subjects that correctly guessed the order was compared to the proportion expected by chance, 1/6, using a one-sample *t*-test (α = 0.05). Note that this is a fairly conservative test, since participants will have the maximum information at the last session to make an educated guess about the condition order; their guess at the end of the study may differ from their perceptions during the study. If we were to find that more people guessed the order correctly than expected by chance, then we planned to test for the effects of guessing correctly similarly to the effects of other covariates. Specifically, in the mixed-effects model examining the effects of L-dopa, we would enter “Guessed Correctly” as a binary covariate, which reflects whether or not the participant accurately guessed that they had received L-dopa on the day that they received it. In the mixed-effects model examining the effects of escitalopram, this covariate would reflect whether or not the participant accurately guessed that they had received escitalopram on the day that they received it.

If there is no evidence that participants guessed the exact medication order correctly, it is still possible that they would be above chance at guessing that they received *some* medication relative to placebo. To test this, we looked again at the order that participants guessed at the end of the study, but instead of counting the order as correct only if all three conditions were properly attributed, we counted the order as correct if the placebo condition was attributed to the right day (even if the escitalopram and L-dopa conditions were switched). This raised the probability of chance performance to 1/3. The proportion of subjects that correctly guessed when they got “something” vs. placebo was compared to 1/3 using a one-sample *t*-test (α = 0.05). If this test revealed above-chance performance, then the mixed-effects models described above would be conducted with “Guessed Placebo Correctly” as a covariate. This covariate reflects whether or not the participant accurately guessed when they were in the placebo condition. Note that either “Guessed Medication Order Correctly,” or “Guessed Placebo Correctly” could be included as a covariate in the primary analyses, but not both (Supplementary Table 1).

##### Effects of medication on affective state

Finally, we did not expect to see significant changes in positive and/or negative affect or state anxiety after either medication. To test this, we performed the following mixed-effects regressions (with random intercepts and coefficients on the medication condition factor by subject) to examine the main effects and interaction effects of medication and time point (before / after) on mood (PANAS positive and negative affect subscales) and anxiety (STAI-S):

STAI-S ∼ 1 + medication * time point + (1 | subject) + (1 | subject: medication)

PANAS (positive affect) ∼ 1 + medication * time point + (1 | subject) + (1 | subject: medication)

PANAS (negative affect) ∼ 1 + medication * time point + (1 | subject) + (1 | subject: medication)

Each regression was conducted twice, once on the L-dopa and placebo data, and once on the escitalopram and placebo data. If we were to find any effects here, then we planned to enter the difference in affect subscale score(s) between the first time point and the second time point as a covariate in the relevant analyses (Supplementary Table 1).

#### Exploratory: Secondary analyses of potential mechanism

We planned to perform the following analyses contingent on finding support for either or both of our Hypotheses (see Supplementary Fig. 4 for flowchart). The purpose of the following analyses was to investigate the mechanism by which L-dopa decreases persistence and/or escitalopram increases persistence. Since these were exploratory analyses, the α level was set at 0.025, the same α level we used for our primary analyses. Note that if we were to find support for Hypothesis 1 *only*, then the following mechanism tests would be restricted to the L-dopa vs. placebo data; if we were to find support for Hypothesis 2 only, then the following mechanism tests would be restricted to the escitalopram vs. placebo data.

*Foraging task.* The foraging task data were examined using linear mixed-effects regression analyses: Leaving time ∼ 1 + block order + travel time * medication + (1 | subject) + (1 | subject: medication) + (1 | subject: medication: travel time)

Here “travel time * medication” encompasses the main effects of travel time and medication, as well as the interaction between them. Here *leaving time* is the average number of harvests per trial in each block, *travel time* is the travel time in that block (two levels: 3 s and 10 s), and *block order* is a categorical factor variable controlling for which travel time condition was presented first (Long first / Short first). Block order was included as a covariate because of evidence that behavior from one foraging environment can carry over to the next^118^. We tested for a main effect of medication condition on leaving time. If the mechanism underlying the effects of our medications on persistence involves perceptions of opportunity cost, then we would expect that, compared to placebo, the leaving time would be shorter in the L-dopa session (indicating more exploration), and the leaving time would be longer in the escitalopram session (indicating more exploitation). In the case that these models are unable to converge with random effects for the interaction terms, we planned to drop random effects terms step-wise until the model did converge.

As detailed in Supplementary Fig. 4, if there is support for our primary hypotheses *and* a significant effect of medication on leaving time in the above foraging model, then we planned to conduct a Sobel mediation test^119,120^ to test for the presence of an indirect effect (specifically, that the medication influences persistence through its effects on foraging). The following regression would be used to test for mediation:

AUC ∼ 1 + leaving time + medication * covariate + (1 | subject) + (1 | subject: medication) + (1 | subject: medication: covariate).

Here “covariate” refers to the covariate(s) that end up being included in the main model. If the coefficient on “medication” is significantly reduced according to the Sobel test statistic, then that suggests that the effects of medication on leaving time in the foraging task are driving the effects of medication on persistence.

For each of the three remaining tasks, the series of analyses conducted would be the same. The dependent variable for the time production task was the accuracy coefficient theta (larger values are indicative of overestimation of time intervals). For the temporal discounting task, the dependent variable was the log-transformed discount rate *k* (higher values indicate more impatience). For the risk tolerance task, the dependent variable was the log-transformed risk tolerance parameter α (higher values indicate more risk tolerance).

We planned to run the following mixed-effects regressions twice, once to examine the effects of L-dopa relative to placebo, and once to examine the effects of escitalopram relative to placebo: Time production accuracy coefficient ∼ 1 + medication + (1 | subject) + (1 | subject: medication) Temporal discount rate ∼ 1 + medication + (1 | subject) + (1 | subject: medication)

Risk tolerance parameter ∼ 1 + medication + (1 | subject) + (1 | subject: medication)

If we were to find no significant effects of medication (at α level of 0.025), then no further tests would be conducted. If there were a difference between the L-dopa and placebo conditions on the control task variable, then we planned to conduct the mediation analysis as described above. The behavior in the control task would be entered as an additional regressor into the regression predicting AUC in the persistence task. For example, if there were an effect of L-dopa on time perception, then the accuracy coefficient would be included in the regression for Hypothesis 1 to test for mediation. If we were to find that the effect of medication on AUC is significantly reduced after controlling for the control task variable, then it is possible that the main behavioral result (e.g., that L-dopa decreases persistence) is driven by the process measured in the control task. The same procedure was also followed for escitalopram, if we were to find any differences between the escitalopram condition and placebo condition on the time production, temporal discounting, or risk tolerance tasks.

## Data availability

All raw data has been shared publicly at this link: https://osf.io/x5ckf/files/osfstorage

## Code availability

All task and analysis code has been shared publicly at this link: https://osf.io/x5ckf/files/osfstorage

## Protocol registration

The registered report protocol (Stage 1 manuscript) can be found at this link: https://springernature.figshare.com/articles/journal_contribution/Effects_of_dopamine_and_serotonin_on_persistence_for_delayed_rewards_in_humans_Registered_Report_Stage_1_Protocol_/19596664

## Acknowledgements

The study was supported by the National Center for Research Resources and the National Center for Advancing Translational Sciences, National Institutes of Health, through Grant UL1TR001878. The funders had no role in study design. The authors would like to thank Ilyssa Delos Reyes and Sage Leland for assistance with recruitment and data collection, and Martin Wiener for advice on the time production task. The authors would also like to thank the University of Pennsylvania’s Investigational Drug Service and Center for Human Phenomic Science.

## Author contributions

All authors were involved in project planning. LZ collected data. KML and LZ cleaned and analyzed data. KML drafted the manuscript, with feedback from AR, DHW, and JWK. All authors reviewed and edited the manuscript.

