## Supplementary Information for "Effects of dopamine and serotonin on persistence for delayed rewards in humans"

**Supplementary Table 1.** Contingency plan for inclusion of covariates

| Covariate | Test | Possible outcomes | Contingency plan |
|---|---|---|---|
| **Variables that may influence persistence or interact with effect of medication on persistence** | | | |
| Body mass index (BMI) | AUC ~ 1 + medication * covariate + (1 \| subject) + (1 \| subject: medication) + (1 \| subject: medication: covariate)<br><br>Here medication refers to: Levodopa vs. Placebo for test of Hypothesis 1<br><br>Escitalopram vs. Placebo for test for Hypothesis 2<br><br>Each covariate in column on the left will be entered one at a time in the analysis above as the "Covariate." | p<0.05 for BMI effect and/or medication*BMI effect | Include BMI in model |
| | | p>0.05 for both BMI effect & medication*BMI effect | Do not include BMI |
| Age | | p<0.05 for age effect and/or medication*age effect | Include age in model |
| | | p>0.05 for both age effect & medication*age effect | Do not include age |
| Sex (Male / Female) | | p<0.05 for sex effect and/or medication*sex effect | Include sex in model |
| | | p>0.05 for both sex effect & medication*sex effect | Do not include sex |
| Condition order (Placebo first / Medication first) | | p<0.05 for order effect and/or medication*order effect | Include condition order in model |
| | | p>0.05 for both order effect & medication*order effect | Do not include condition order |
| Working memory (WM) capacity (operation span): **For Hypothesis 1 only** | | p<0.05 for WM effect and/or medication*WM effect | Include WM in model |
| | | p>0.05 for both WM effect & medication*WM effect | Do not include WM |
| Barratt Impulsiveness Scale (BIS total): **For Hypothesis 1 only** | | p<0.05 for BIS effect and/or medication*BIS effect | Include BIS in model |
| | | p>0.05 for both BIS effect & medication*BIS effect | Do not include BIS |
| **Assessing effectiveness of blinding** | | | |
| Step 1. "Guessed medication order correctly" | Proportion of subjects that correctly guessed the medication order will be compared to chance, 1/6, using a one-sample *t*-test | p<0.05 in one-sample *t*-test | Include "Guessed correctly" in model |
| | | p>0.05 in one-sample *t*-test | Proceed to Step 2 |
| Step 2. "Guessed placebo correctly" | Proportion of subjects that correctly guessed when they got an active drug vs. placebo will be compared to chance, 1/3, using a one-sample *t*-test | p<0.05 in one-sample *t*-test | Include "Guessed placebo correctly" in model |
| | | p>0.05 in one-sample *t*-test | Blinding was successful; do not include either indicator as a covariate |
| **Changes in affect as a result of pharmacological manipulation** | | | |
| PANAS-N (Negative affect subscale) | PANAS-N ~ 1 + medication * time point + (1 \| subject) + (1 \| subject: medication)<br>Where *time point* refers to before or after manipulation. | p<0.05 for main effect of medication and/or medication*time effect | Include Change in PANAS-N (Time 2 minus Time 1) as covariate in model |
| | | p>0.05 for both main effect of medication and medication*time effect | Do not include PANAS-N |
| PANAS-P (Positive affect subscale) | PANAS-P ~ 1 + medication * time point + (1 \| subject) + (1 \| subject: medication) | p<0.05 for main effect of medication and/or medication*time effect | Include Change in PANAS-P (Time 2 minus Time 1) as covariate in model |
| | | p>0.05 for both main effect of medication and medication*time effect | Do not include PANAS-P |
| STAI-S (State anxiety) | STAI-S ~ 1 + medication * time point + (1 \| subject) + (1 \| subject: medication) | p<0.05 for main effect of medication and/or medication*time effect | Include Change in STAI-S (Time 2 minus Time 1) as covariate in model |
| | | p>0.05 for both main effect of medication and medication*time effect | Do not include STAI-S |

**Supplementary Table 2.** Fixed effects of SSRI on leaving threshold in foraging task.

| **Predictor** | **β** | **95% CI** |
|---|---|---|
| SSRI<br>(1 = SSRI; 0 = Placebo) | -0.47 | -0.97, 0.04 |
| Travel Time Block<br>(1 = Long; 0 = Short) | -1.42*** | -1.82, -1.02 |
| SSRI x Travel Time Block | -0.07 | -0.50, 0.36 |
| Block Order<br>(1 = Long first; 0 = Short first) | -0.78** | -1.35, -0.21 |
| SSRI x Block Order | 0.46 | -0.25, 1.18 |
| Travel Time Block x Block Order | 0.89** | 0.32, 1.46 |
| SSRI x Travel Time Block x Block Order | -0.16 | -0.77, 0.45 |

Note: Dependent variable is leaving threshold (average number of berries at last harvest) from foraging task, with higher values indicating leaving earlier. Random effects were included for the intercept, SSRI, Travel Time Block, and Block Order terms. CI = confidence interval. SSRI = selective serotonin reuptake inhibitor (20 mg escitalopram). **$p<0.01$, ***$p<0.001$.

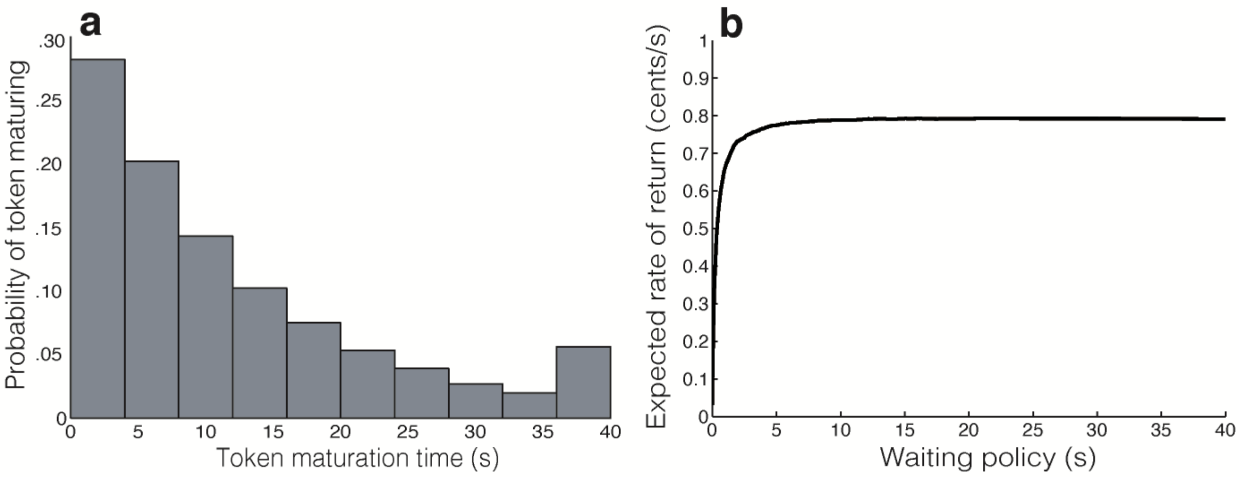


**Supplementary Fig. 1.** Reward timing distribution in persistence task, based on simulation (100,000 samples). Reward (i.e., token maturation) probability as a function of elapsed time (a), and projected (average) earnings per block as a function of waiting policy (b).

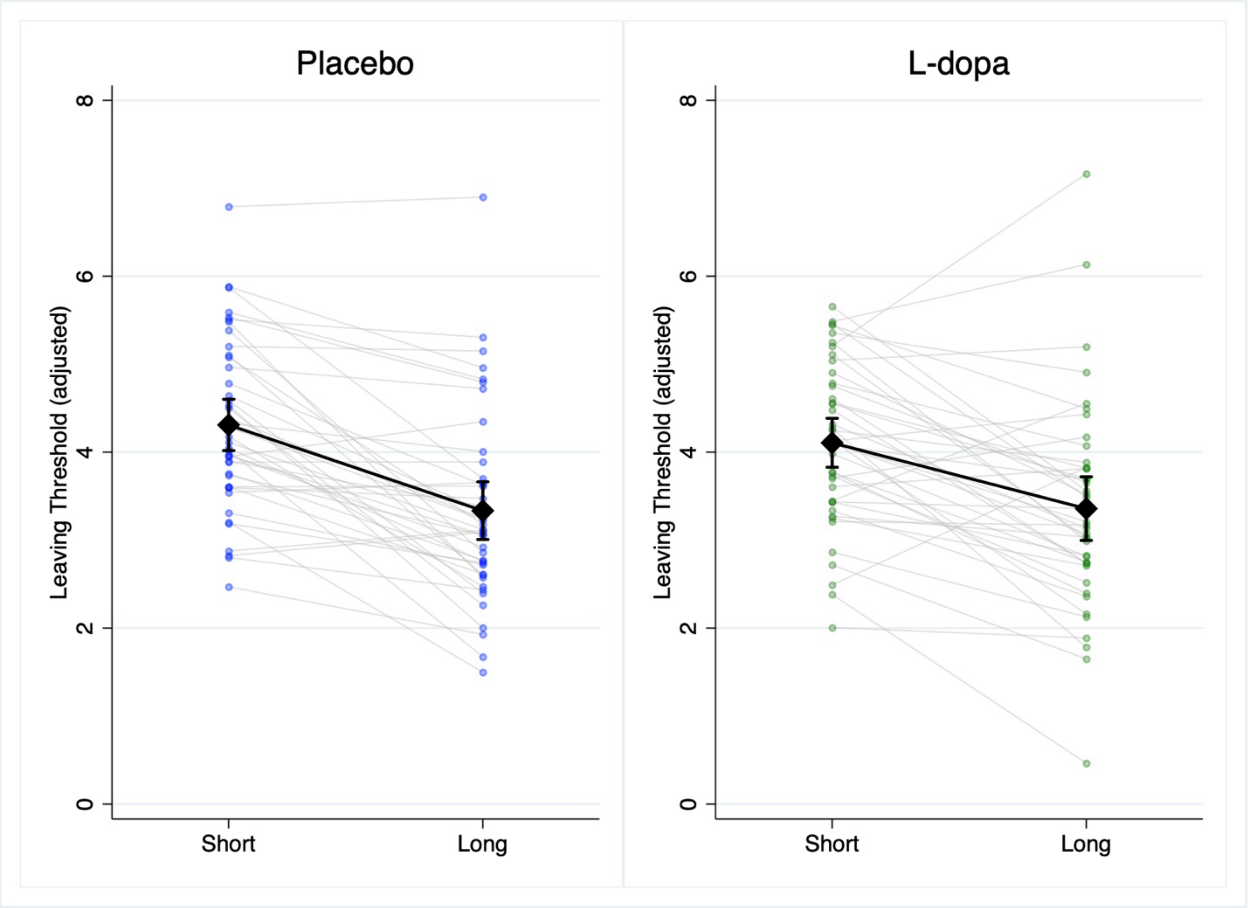


**Supplementary Fig. 2.** Leaving threshold data from foraging task for placebo and L-dopa conditions, adjusted for foraging order. Participants are represented by dots, with connectors showing change across conditions. The panel on the left shows data from the placebo condition, and the panel on the right shows data from the L-Dopa condition. Means and 95% confidence intervals are also shown. Short = short travel time block. Long = long travel time block. Whereas the leaving threshold was higher in the short travel time condition overall (participants left patches earlier when opportunity cost was higher), there was no effect of L-dopa on the overall leaving threshold. However, there was an interaction between L-dopa and travel time on leaving threshold in our mixed-effects model controlling for foraging order, such that L-dopa reduced the influence of opportunity cost on leaving threshold (β = 0.45; 95% CI [0.02, 0.87]; $p$ = 0.039).

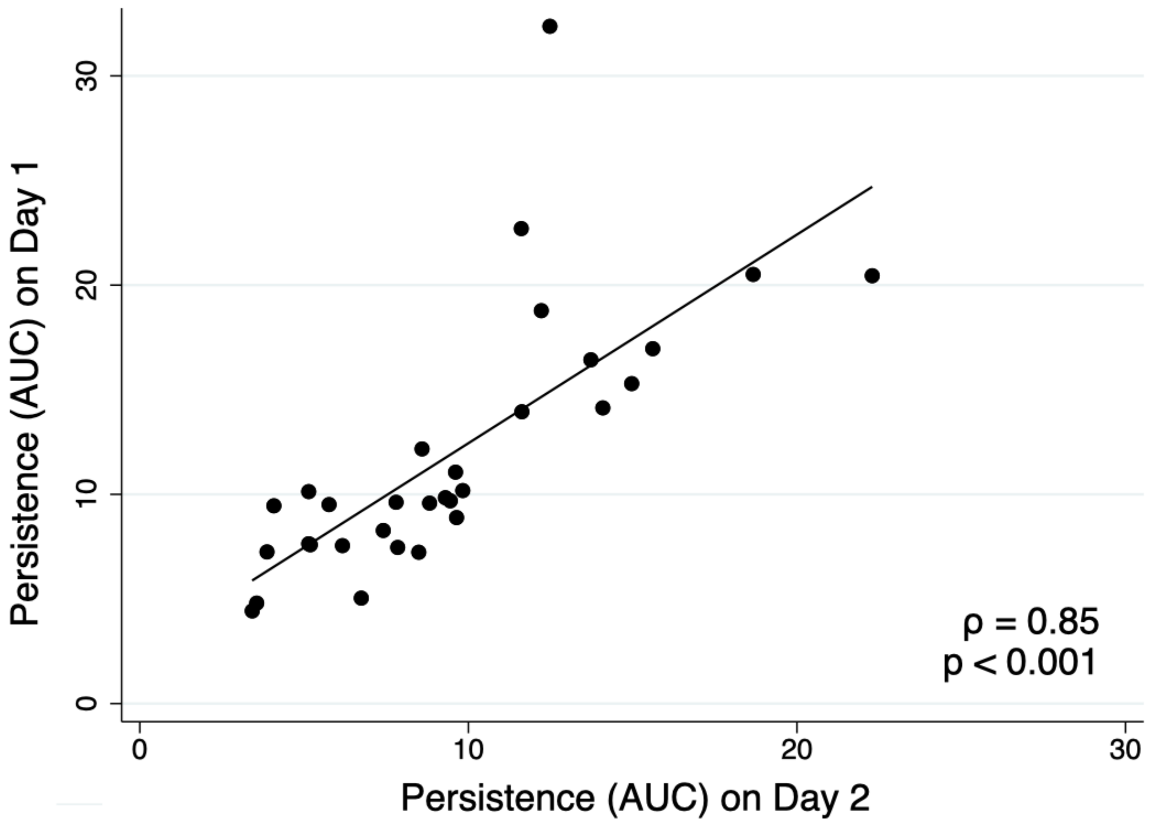


**Supplementary Fig. 3.** Pilot data from our persistence task. Persistence is test-retest reliable over one week (Spearman $\rho = 0.85$; $p < 0.001$; $n = 31$).

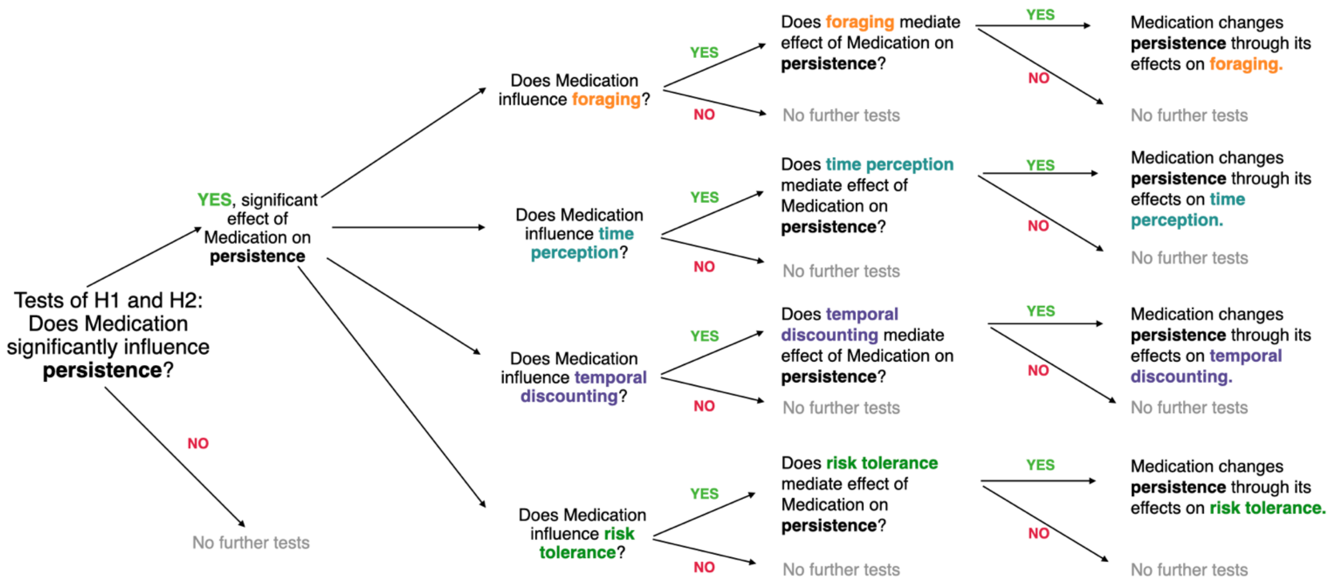


**Supplementary Fig. 4.** Exploratory analyses of mechanism. The whole process summarized above was done separately for our two hypotheses. If we were to find support for the hypothesis (H1 or H2), we then planned to test to see whether the medication (levodopa for H1; escitalopram for H2) influenced behavior in any of the potential mechanism tasks (Foraging, Time Perception, Temporal Discounting, and Risk Tolerance). If so, then we planned to conduct Sobel tests of mediation to determine whether there is a significant indirect effect whereby the medication influences persistence via the mechanism measured in that task. The results of those tests would shed light on whether any of the secondary tasks mediate the relationship between medication status and persistence.
